# High-throughput behavioural phenotyping of 25 *C. elegans* disease models including patient-specific mutations

**DOI:** 10.1101/2025.02.07.637026

**Authors:** Thomas J. O’Brien, Eneko P. Navarro, Consuelo Barroso, Lara Menzies, Enrique Martinez-Perez, David Carling, André E. X. Brown

## Abstract

Genetic diagnosis is fast and cheap, challenging our capacity to evaluate the functional impact of novel disease-causing variants or identify potential therapeutics. Model organisms including *C. elegans* present the possibility of systematically modelling genetic diseases, yet robust, high-throughput methods have been lacking. Here we show that automated multi-dimensional behaviour tracking can detect phenotypes in 25 new *C. elegans* disease models spanning homozygous loss-of-function alleles and patient-specific single-amino-acid substitutions. We find that homozygous loss-of-function (LoF) mutants across diverse genetic pathways (including BORC, FLCN, and FNIP-2) exhibit strong, readily detectable abnormalities in posture, locomotion, and stimulus responses compared to wild-type animals. An *smc-3* mutant strain—modelled by introducing a patient-identified missense change—causes developmental anomalies and distinct behavioural profiles even though complete loss of SMC-3 is lethal. In contrast, patient-derived missense mutations in another essential gene, *tnpo-2*, did not show a strong phenotype initially but it could be “sensitized” chemically (e.g., with aldicarb), potentially facilitating future drug screens. Our findings show that scalable behavioural phenotyping can capture a wide range of mutant effects—from strong to subtle—in patient-avatar worm lines. We anticipate that this standardized approach will enable systematic drug repurposing for rare genetic disorders as new disease variants are discovered.

## Introduction

Advances in genome and exome sequencing technologies mean the rates of genetic variant classification and drug development cannot keep pace with the rate of disease gene discovery. More than 100 genes associated with new genetic disorders are discovered and uploaded to databases such as OMIM^1,2^ every year. Genetic diagnosis of disease is an important first step, but identifying the genetic cause associated with the onset of a disorder does not necessarily lead to a drug target hypothesis. This is particularly true for rare diseases which are often poorly characterised and underfunded: ∼95% have no approved treatment or validated therapeutic target^2^. Approximately 74% of rare diseases affect the central nervous system^3^, giving rise to pleiotropic presentations of disease in both the clinic and *in vivo* vertebrate models, which complicates investigation into disease onset and progression^4,5^. Moreover, different variants within the same gene can result in different phenotypes^6^. There is therefore a need to develop methods to efficiently study patient-specific mutations to support the development of personalised therapies.

In the absence of a validated drug target, phenotypic screens in whole organism models provide an alternative route for drug discovery. However, the use of *in vivo* models in high-throughput studies can be hampered by the lack of a robust and rapidly detected phenotype. The cost and speed of making and validating new genetic and phenotyping methods for new disease/allelic variants is a further challenge.

We have recently shown that high-throughput tracking shows promise for systematic phenotyping and drug repurposing using a single multiwell plate assay^7^. Using *C. elegans* as a model organism, we followed a disease-phenolog approach to develop a standardised screening method that combines high-resolution worm tracking^8^ with automated quantitative behavioural phenotyping^7,9–14^. Because we detect multiple phenotypes with a single assay, multiple mutations or treatment conditions can be screened using the same approach without prior knowledge of the molecular underpinnings of a disease other than the mutation to model.

Previously we focussed on diseases that could be modelled using knockouts. Here we extend the approach to characterise 25 additional *C. elegans* disease models encompassing homozygous loss-of-function (LoF), heterozygous LoF, and single amino acid substitution mutants. We detected behavioural differences for all strains compared to wild-type. The mutations affect diverse genetic pathways and we detected correspondingly diverse phenotypes. Despite this diversity, we find that mutations in genes that are involved in related cellular processes lead to similar phenotypic profiles.

The multidimensional behavioural fingerprinting of *C. elegans* patient avatars provides a foundation for scalable drug discovery efforts using a readily-perturbable whole organism model. The systematic creation, phenotyping, and screening of patient-specific genetic variants may form the basis for the discovery of lead compounds and the development of personalised therapeutics at a rate commensurate with disease gene discovery.

## Results

### Behavioural phenotypes are detected for all disease model mutants

The 25 mutant strains made for this study are primarily based on genetic variants found in patients that were identified by collaborators and they cover 25 different genes associated with a diverse representation of rare genetic disorders. Different strategies were used depending on the specific mutation and its phenotype. These are detailed in the strain-specific gene cards (supplementary information). All mutants were initially characterised using a previously developed pipeline^7,8,11–14^. Briefly, young adult worms are added to the wells of a 96-well plate using a COPAS Biosort and recorded on custom imaging rigs (LoopBio). We record a 16-min video comprising: (1) a 5-min pre-stimulus recording (capturing differences in baseline behaviour), (2) a 6-min blue light recording (testing differences in a strains’ photophobic escape response^15^), (3) a 5-min post-stimulus recording (testing recovery to stimulation). We then use Tierpsy^9^ to extract 2763 features (covering morphology, posture and locomotion) for each video which are concatenated to make a phenotypic profile which is averaged over the worms in each well (8289 features in total)^10,16^.

Hierarchical clustering of the different patient avatars reveals that different behavioural phenotypes are captured across this diverse panel of mutant strains (Figure 1A). Phenotypic similarities are also captured for strains that have mutations within genes predicted to have a similar function or which are associated with similar diseases in humans. For example, *blos-1*, *blos-9*, and *sam-4* are all predicted to be members of the BLOC-one-related complex (BORC)^17^. LoF mutations within these genes result in very similar behavioural phenotypes. However, this is not always the case, as our *blos-8* LoF mutant (also predicted to be a subunit of the BORCS complex) has a markedly different behavioural phenotype.

**Figure 1.**
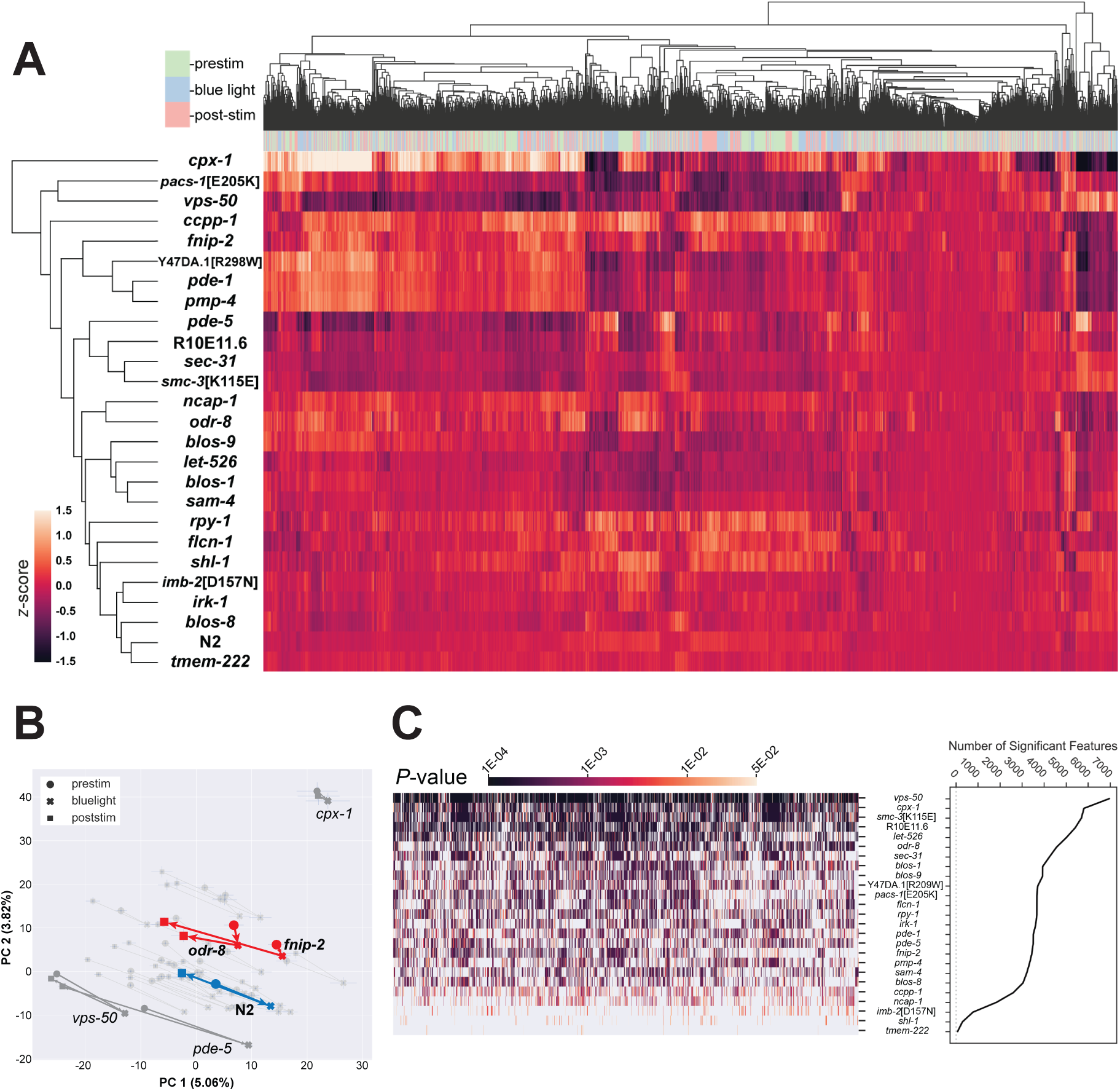
All disease model mutants exhibit behavioural differences compared to the control. (A) Hierarchical clustering of the entire behavioural feature set extracted by Tierpsy (8289 features total). Features are *Z*-normalised. The top barcode shows when during the period of image acquisition the behavioural features were extracted: pre-stimulation (pink), blue light (blue) and post-stimulation (green). (B) Principal component analysis of the disease model mutants and wild-type reference (blue). Strains move in phenospace between pre-stimulus (circular points), blue light (crosses) and post-stimulation (squares) recordings. Mutant strains that do not return towards their pre-stimulus position in phenomic space are coloured red. (C) The total number of statistically significant behavioural features extracted for each strain compared to N2. *P*-values for each feature are calculated using block permutation t-tests, using *n* = 100,000 permutations and *P* < 0.05 is considered statistically significant after correcting for multiple comparisons using the Benjamini-Yekutieli method.

We next performed principal component analysis (PCA) on the same data, but separated into the pre-stimulus, blue light, and post-stimulus tracking (Figure 1B). Blue light stimulation affects worm behaviour which corresponds to a shift of points in phenotype space. Most strains show a partial phenotypic recovery (similar to wild-type N2) during post-stimulation recordings. However, *odr-8*(*syb4940*) and *fnip-2*(*syb8038*) mutants show a reduced recovery (Figure 1B).

Of the 25 disease model strains, 22 had a ‘strong’ behavioural phenotype, exhibiting >1000 statistically significant behavioural differences compared to N2 (Figure 1C). These strains are more likely to be suitable for drug screens where many compounds are tested, which would typically necessitate a smaller number of replicates. Strain-specific phenotype summaries, the molecular characteristics and human disease(s) associated with each strain are available in the supplementary information (strain-specific gene cards). We have also created an interactive heatmap of the hierarchical cluster map (Figure 1A) that can be used to explore the behavioural differences of all strains compared to wild-type (available at https://zenodo.org/records/13941390).

### BORC complex mutations

The BLOC-one-related complex (BORC) is a multi-subunit protein complex that enables the correct positioning of lysosomes within cells^17^. By promoting the recruitment of the small GTPase ARL8^18^, which in-turn recruits the motor proteins kinesin-1^19^ and kinesin-3^20^, BORC enables the anterograde (centrifugal) movement of lysosomes along microtubule tracks in non-polarised cells^17,19,21^ and enables movement towards the distal axon tip in neurons^22–24^. Truncation of any member of the complex results in the juxtanuclear positioning of lysosomes^25^. Furthermore, genetic variants within BORC genes are associated with many neurodegenerative disorders^26^ including: Hermansky-Pudlak Syndrome^27^, hereditary spastic paraplegia^24,28,29^, Parkinson’s Disease^30,31^, Huntingdon’s Disease^32^, Alzheimer’s^33^, amyotrophic lateral sclerosis^34^, schizophrenia^35,36^, and neuronal axonal dystrophy^37^. Despite its widespread expression, the consequences of BORC deficiency primarily manifest in the nervous system. Thus, BORC is an emerging research target for the treatment of many neurodevelopmental conditions^30,38^.

BORC is conserved between vertebrate^6^, invertebrate^20,39–41^ and prokaryotic^42^ species. *C. elegans* embryos have been used to study defective phagolysosome clearance in *sam-4* (*BORCS5* ortholog) and *blos-7* (*BORCS6* ortholog) LoF mutants^41^, and *borcs8* knockout (*BORCS8* ortholog) zebrafish are found to have general neurodevelopmental delay and locomotive defects^6^. Here we selected 4 genes associated with BORC: *blos-1* (*BLOCS1* ortholog), *blos-8* (*BORCS7* ortholog), *blos-9* (*BORCS9* ortholog), and *sam-4* and created homozygous deletions to model BORC deficiency.

Mutants with homozygous deletions of these BORC genes are viable (in contrast to vertebrates^6,22,24^) and show strong behavioural phenotypes with >3000 features significantly different from N2. *blos-1*(*syb6895*), *blos-9*(*syb7029*) and *sam-4*(*syb6765*) are all shorter, whereas *blos-8*(*syb6686*) is longer, than N2 (Figure 2A). Furthermore, *blos-1*, *blos-9*, and *sam-4* LoF primarily affects head-related features, including decreased angular velocity (Figure 2B), curvature (Figure 2C), and acceleration or speed of the head (strain-specific gene cards, Supplementary Information). However, *blos-8* LoF caused a significant increase in head curvature (Figure 2C) and affected other areas of the worm body, for example resulting in a decrease in tail angular acceleration (Figure 2D). We detect no differences in the fraction of time worms spend moving forwards between wild-type (N2) worms and *blos-8*(*syb6686*) or *sam-4*(*syb6765*), but *blos-1*(*syb6895*) has an increased forward and decreased backward response to stimulation with blue light (Figure 2E-G). In contrast, *blos-9*(*syb7029*) has a decreased forward and increased backward response.

**Figure 2.**
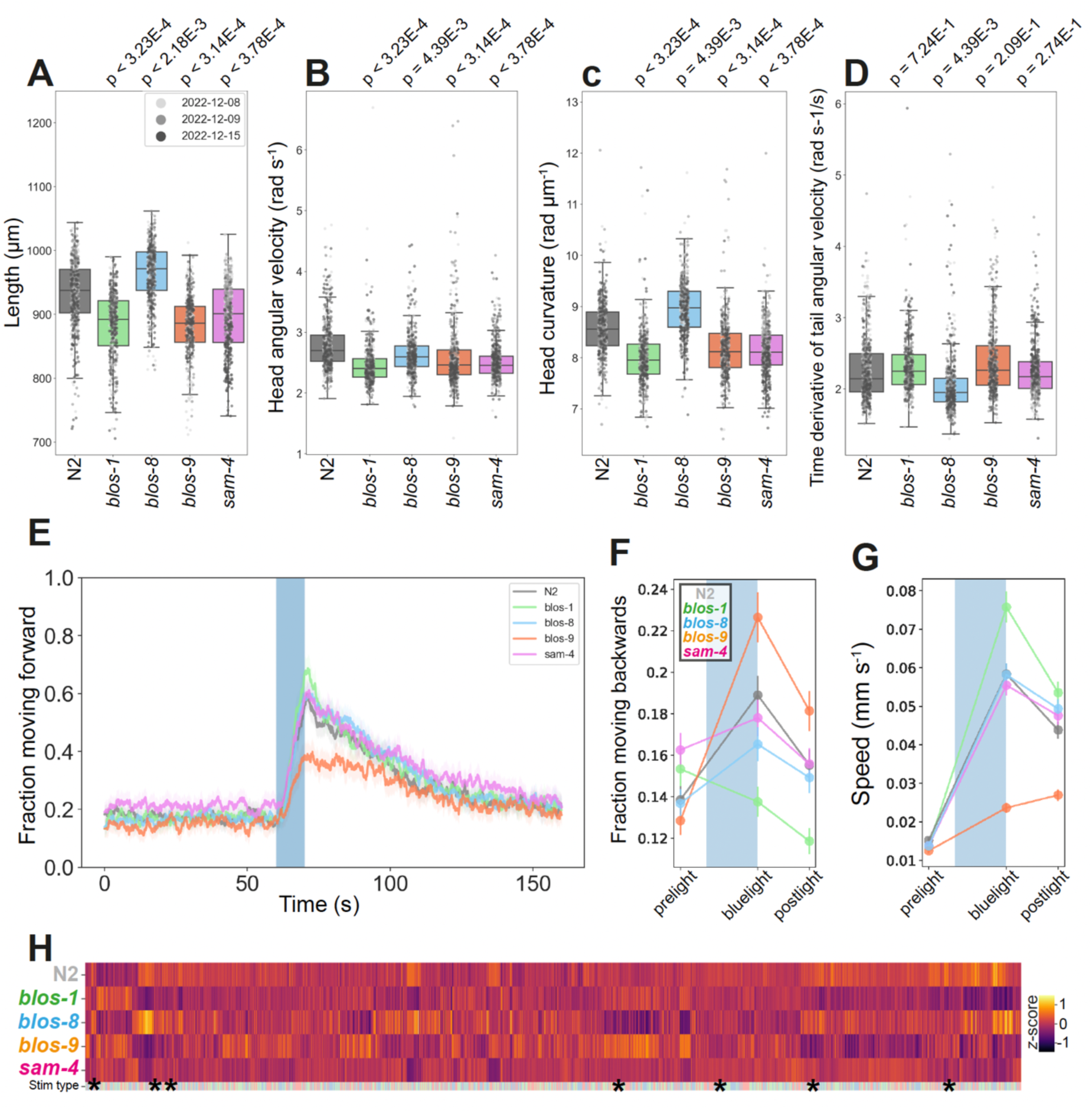
BORC deficiency disease model phenologs. (A-D) Key behavioural features altered in strains containing loss-of-function mutations in orthologs associated with BORC: *blos-1*(*syb6895*), *blos-8*(*syb6686*), *blos-9*(*syb7029*) and *sam-4*(*syb6765*). Individual points marked on the box plots are well averaged values (3 – 5 worms per well) for each feature across the independent days of tracking. *P*-values are for comparison to wild-type N2 using block permutation t-tests (*n* = 100,000 permutations, correcting for multiple comparisons using the Benjamini-Yekutieli method). (E) Overall fraction of worms moving forward 60 s prior to and 80 s following stimulation with a 10 s blue light pulse (blue shading). Coloured lines represent averages of the detected fraction of paused worms across all biological replicates and shaded areas represent 95% confidence intervals. (F-G) Changes in selected features in response to stimulation with a single 10 second pulse of blue light (shaded region). Feature values are calculated using 10 second windows centred on 5 seconds before, 10 seconds after, and 20 seconds after the beginning of the pulse. (H) Heatmap of the entire set of 8289 behavioural features extracted by Tierpsy for the disease model strains associated with BORC deficiency. The ‘stim type’ barcode denotes when during image acquisition a feature was extracted: pre-stimulation (pink), blue light stimulation (blue) and post-stimulation (green). Asterisks show the location of the selected features presented in A-D.

### Folliculin mutations

Birt-Hogg-Dubé (BHD) syndrome is a rare autosomal dominant disorder characterised by fibrofolliculomas, lung cysts, spontaneous pneumothorax and renal cell carcinomas^43,44^. Variants in the *FLCN* gene are found to be responsible for BHD^43^. FLCN is highly conserved from unicellular organisms to mammals^45^. Studies in mice identified two conserved binding partners of FLCN, FNIP1 and FNIP2, that regulate AMPK (5′-AMP activated protein kinase) activation^46,47^. Despite this known interaction, their precise role is unclear as both inhibition and stimulation of AMPK have been reported^45^. Recently an association of FNIP1/FINP2 in glucose homeostasis has been confirmed using a mouse model with an adipose tissue specific ablation of FNIP1^48^. FLCN is also reported to have a role in mTOR suppression^49^, mTORC regulation^50–53^, ciliogenesis^54^, TGF-β signalling^55,56^, autophagy^57–59^, cell adhesion^53^, cell polarity^60^, and regulation of the cell cycle^61,62^.

Homozygous deletion of *flcn-1* (FLCN ortholog) or *fnip-2* (FNIP1 and FNIP2 ortholog) in *C. elegans* is non-lethal, causes no developmental delay and results in strong behavioural phenotypes (3663 or 3433 statistically significant differences, respectively, compared to N2) (Figure 3). *flcn-1*(*syb8071*) is longer than N2 with no difference in width (Figure 3A-B). Whereas *fnip-2*(*syb8038*) is shorter and wider^63^. Both mutants have increased angular velocity for all body segments (Figure 3C) and increased time derivative of curvature (Figure 3E and strain-specific gene cards). Whereas deletion of *flcn-1* causes a significant increase in head curvature, deletion of *fnip-2* results in a decrease of this feature (Figure 3D). Both mutants are hyperactive (Figure 3F), have a decreased backward blue light response (Figure 3G), and a more sustained forward blue light response (Figure 3H).

**Figure 3.**
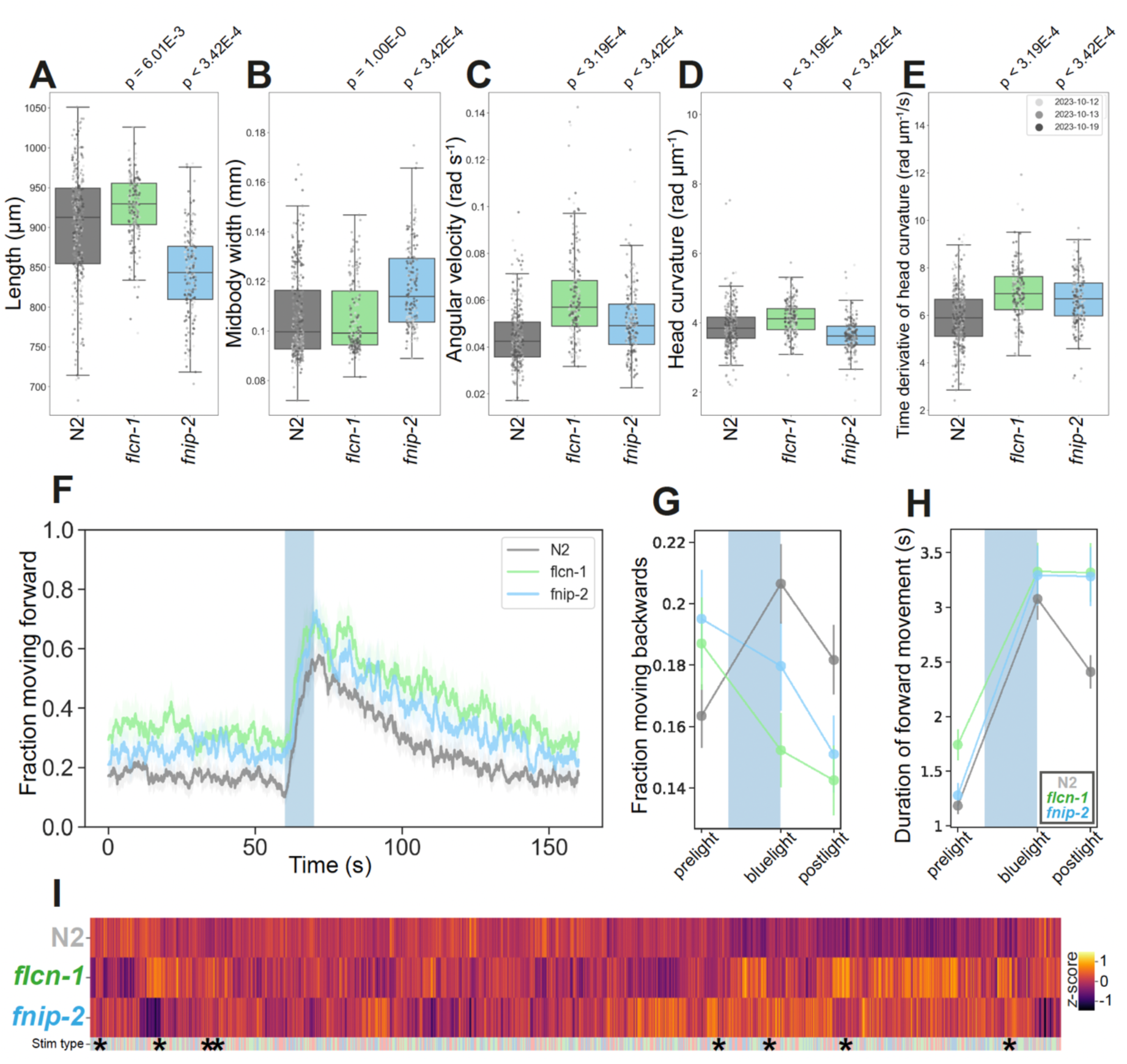
Folliculin mutant disease model phenologs. (A-E) Key behavioural features altered in *s*trains containing loss-of-function mutations in the *C. elegans* orthologs of *FLCN*, *flcn-1*(*syb8071*), or *FNIP1/2*, *fnip-2*(*syb8038*). Individual points marked on the box plots are well averaged values (3 – 5 worms per well) for each feature across the independent days of tracking. *P*-values are for comparison to wild-type N2 using block permutation t-tests (*n* = 100,000 permutations, correcting for multiple comparisons using the Benjamini-Yekutieli method). (F) Overall fraction of worms moving forward 60 s prior to and 80 s following stimulation with a 10 s blue light pulse (blue shading). Coloured lines represent averages of the detected fraction of paused worms across all biological replicates and shaded areas represent 95% confidence intervals. (G-H) Changes in selected features in response to stimulation with a single 10 second pulse of blue light (shaded region). Feature values are calculated using 10 second windows centred on 5 seconds before, 10 seconds after, and 20 seconds after the beginning of the pulse. (I) Heatmap of the entire set of 8289 behavioural features extracted by Tierpsy for our *flcn-1* and *fnip-2* LoF mutants. The ‘stim type’ barcode denotes when during image acquisition a feature was extracted: pre-stimulation (pink), blue light stimulation (blue) and post-stimulation (green). Asterisks show the location of the selected features presented in A-E

Given the role of FLCN and FNIP1/2 in the regulation of AMPK and mTOR signalling, we tested a panel of compounds known to modulate AMPK/mTOR pathways to determine how they altered the behaviour of *flcn-1*(*syb8071*), *fnip-2*(*syb8038*) or N2. Due to the known antimicrobial activity of rapamycin (an mTOR inhibitor), we used PFA-killed OP50 as a food source for these experiments^64^.

Pairwise comparison of feature vectors extracted from strains exposed to DMSO only (untreated) vs 4h treatment with 100μM of each compound revealed no discernible change in the behaviour of any strain treated with rapamycin (Figure 4A). Treatment with 3BDO (mTOR activator) has no effect on N2 or *fnip-2*(*syb8038*), but leads to 249 significant feature differences between untreated and treated *flcn-1*(*syb8071*) mutants (Figure 4A-B). Although this is a weak phenotype, there is a consistent change in some speed-related features, alongside rescue of the decreased speed phenotype we observed between untreated N2 and the untreated *flcn-1* LoF strain (Figure 4B).

**Figure 4.**
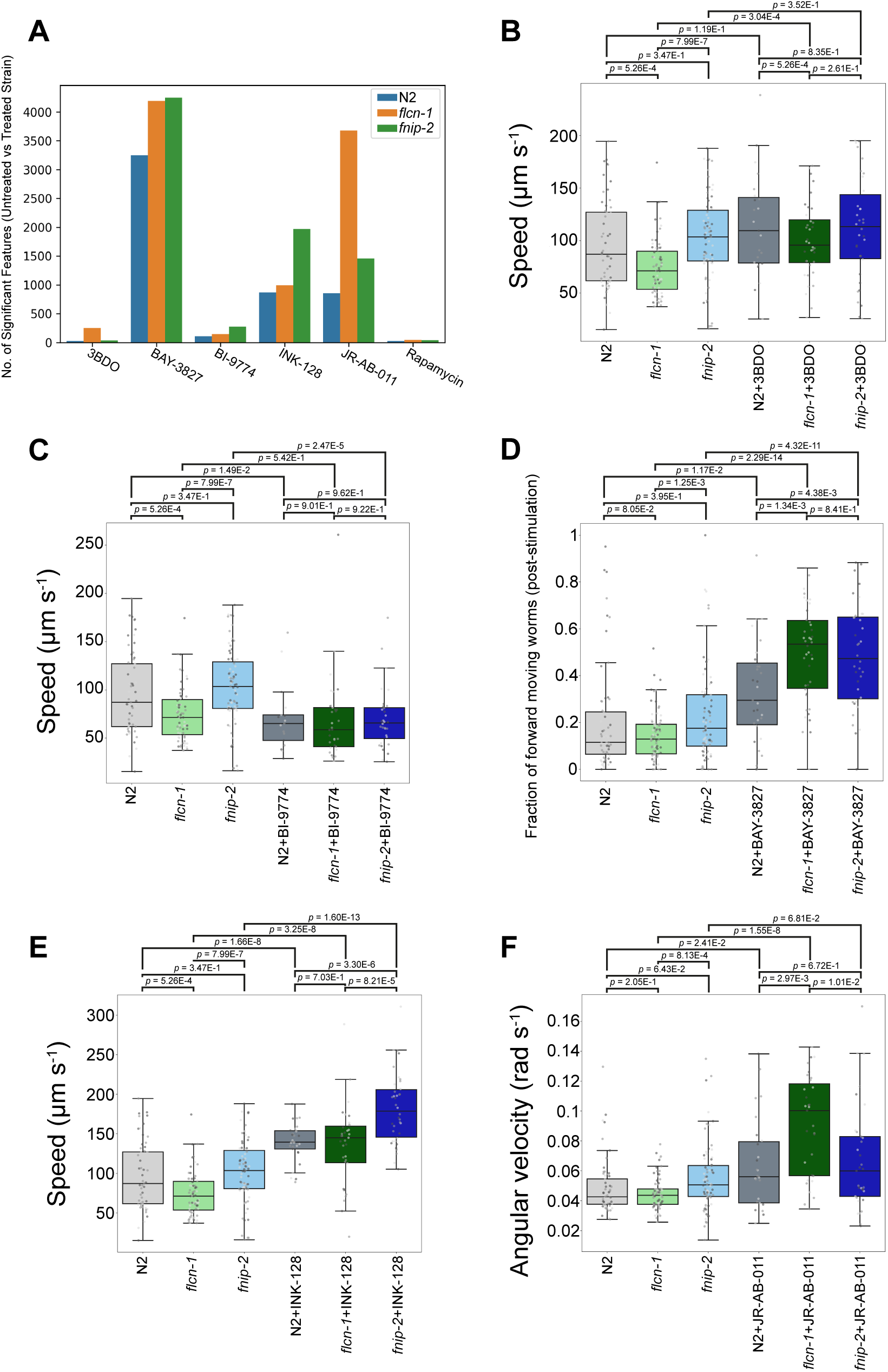
Effect of AMPK and mTOR inhibition or activation of folliculin mutant phenotypes. (A) Total number of behavioural changes detected between either N2, *flcn-1*(*syb8071*) or *fnip-2*(*syb8038*) treated with a compound vs the same strain treated with DMSO (untreated). Worms were exposed to 100 mM of each compound for 4h prior to tracking. Statistically significant phenotypic changes were calculated using the Kruskal-Wallis test correcting for multiple comparisons using the Benjamini-Yekutieli method. (B-F) Key behavioural features differing between untreated strains and worms treated with the various AMPK and mTOR activator or inhibitors. Individual points marked on the box plots are well averaged values (3 – 5 worms per well) for each feature across the independent days of tracking.

Treatment with the pan-AMPK activator, BI-9774, causes a significant decrease in the speed of N2 and *fnip-2*(*syb9038*). There is no statistically significant difference in the speed of any of the treated strains after 4h exposure to this compound (Figure 4C). Although the phenotypic changes observed in *fnip-2*(*syb8038*) as a result of BI-9774 treatment are weak (only 337 significant differences), the differences are consistent with an increased susceptibility to AMPK activation in mutants lacking FNIP1/2. In contrast, treating worms with BAY-3827 (selective AMPK inhibitor) causes a very strong change in the behaviour of the wild-type and mutant strains (Figure 4A), primarily causing an increase in most speed-related features. There are >1000 significant phenotypic changes detected between the BAY-3827 treated and untreated mutants compared to the number of phenotype changes detected between untreated N2 and N2 treated with the BAY-3827 (Figure 4A), suggesting that deletion of *flcn-1* or *fnip-2* makes worms more susceptible to AMPK inhibition. As an example, both mutants exhibit a greater fraction of the population moving after the cessation of blue light stimulation (Figure 4D).

Like AMPK inhibition, treatment with INK-128 (an mTOR1/2 inhibitor) primarily causes an increase in speed-related features across all the strains (Figure 4E). Exposure to INK-128 results in a >2-fold increase in the number of behavioural differences detected between the treated and untreated *fnip-2* LoF worms (1969 feature differences) compared to *flcn-1*(*syb8071*) (994 features) or N2 (873 features, Figure 4A). Furthermore, the increase in speed as a result of mTOR1/2 inhibition is more pronounced for the *fnip-2* LoF mutant compared to the other strains (Figure 4E). Treatment with the selective mTORC2 inhibitor, JR-AB-011, caused a similar increase in the speed and locomotive-related features for all strains and we note little difference in the total number of phenotypic changes between N2 treated with INK-128 or JR-AB-011 (Figure 4A). The *flcn-1* LoF mutant is more susceptible to treatment with JR-AB-011 and >2000 more significant phenotypic differences are observed for treated *flcn-1*(*syb8071*) compared to *fnip-2*(*syb8038*) or wild-type. For example, we observe a strong increase in the angular velocity of the *flcn-1* LoF mutant upon JR-AB-011 treatment that is not observed in the other strains (Figure 4F). Inhibition of mTOR1/2 results in more pronounced phenotypic changes (increased susceptibility) for worms lacking the FNIP1/2 ortholog, but knock-out of the FLCN ortholog results in a greater susceptibility of worms to mTORC2 inhibition.

We also attempted to use Western Blotting to validate the molecular underpinnings of these phenotypic changes as a result of differences in AMPK and mTOR signalling. For this we tried to measure the phosphorylation of T172 in AMPKα or S49 (worm position of human S51) eIF2α as a measure of AMPK^65^ or mTORC1^66,67^ signalling, respectively. Although we observed differences in the basal level of signalling between the untreated strains, we could not reliably identify changes in the phosphorylation state of either site following treatment with the different drugs (Supplementary Information 1). These results could be due to a lack of specificity of the available antibodies to worm AMPKα and eIF2α phosphorylation sites or the transient nature of the phosphorylation states.

### Enhancing the separation of a TNPO2 patient avatar in phenomic space

The behavioural phenotypes we describe above relate to strains containing CRISPR deletions of an entire gene coding region (putative LoF mutants). However, some pathological mutations do not cause complete loss of function so a knockout may not be the best model. We created several strains with patient-specific single amino acid changes: *imb-2[D157N]* (ortholog of human *TNPO2[D156N]*), *pacs-1[E205K]* (ortholog of human *PACS2[E209K]*), *smc-3[K115E]* (ortholog of human *SMC3[K114E]*), and *Y47D91A.1[R298W]* (ortholog of human *GMPPA[R318W]*).

The *pacs-1[E205K]*, *smc-3[K115E]* (described in detail in the next section), and *Y47D91A.1[R298W]* mutants exhibit strong behavioural phenotypes (>3500 significant features compared to wild-type, see strain-specific gene cards), so are well suited for high-throughput drug screens. However, *imb-2[D157N]* exhibits a moderate behavioural phenotype (770 significant features). This was somewhat surprising because the mutation is associated with severe disease in humans and *imb-2* deletion and RNAi are lethal in worms^73–75^. TNPO2 variants cause a number of neuronal abnormalities. Both loss and gain of transportin activity cause developmental defects^76^ and disease severity has been found to depend on the position of mutations within the protein^76,77^.

Although we detect 770 behavioural differences between *imb-2*(*syb6372*) and N2 (strain specific gene card), these are relatively small changes that are detected with a large number of well replicates (*n* > 300). Random sub-sampling of the tracking data to simulate sample sizes typical for a drug screen makes phenotype detection unreliable (Supplementary Figure 2). We therefore tested a number of screening conditions to enhance the separation of these strains in phenomic space to enable a drug screen. *imb-2*(*syb6372*) is homozygous for the aspartic acid to asparagine variant, whereas the corresponding patient variant is heterozygous. Given that heterozygous overdominance is known to impact fitness and that certain diseases only manifest in heterozygous patients, *e.g.,* PCHDH19-related epilepsy^78^, we first determined the behaviour of a heterozygous variant (*imb-2[D157N]/+*). Phenotypic comparison of *imb-2[D157N]/+* and N2/+ revealed no behavioural differences between the heterozygous strains (data not shown), similar to prior findings that *imb-2*(*vy10*) (containing a different single amino acid change) is recessive in *C. elegans*^77^.

Alongside *imb-2*, *C. elegans* encodes *imb-1* (*KPNB1* ortholog) and *imb-3* (*IPO5* and *RANBP6* ortholog) that also play a role in the import of proteins into the nucleus^79–82^. Due to potential functional redundancy, we used RNAi to knock down *imb-1* or *imb-3* in *imb-2* mutants (Figure 5A-C). We fed *imb-2* mutants bacteria expressing dsRNA targeting *imb-1* and *imb-3* as larvae and tracked worms as adults. There was not a large effect of RNAi, but focusing on the subset of features highlighted in the *imb-2* gene card (Supplementary Information), silencing of *imb-1* resulted in the best separation of strains with respect to increased midbody curvature, increased tail speed, and decreased head foraging (time derivative of angular velocity) behaviours (Figure 5A-C). Still, sub-sampling the tracking data to simulate a drug screen with *n* = 3 replicates shows that the phenotype is not robust (Supplementary Figure 2B).

**Figure 5.**
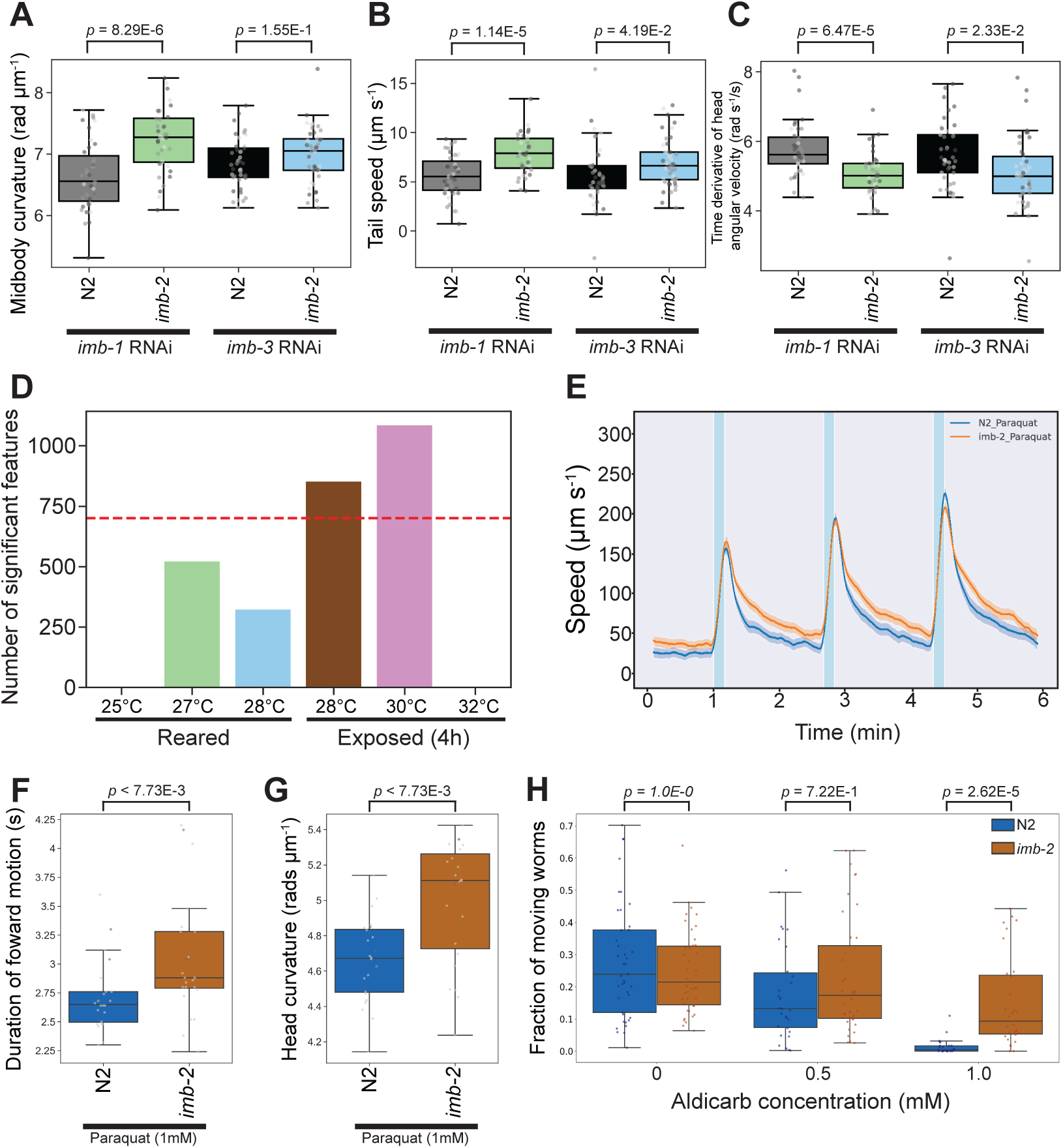
Enhancing the phenomic separation of *TNPO2* mutants and controls. (A-C) Key behavioural differences of *imb-2*(*syb6372*) and wild-type worms following the RNAi-mediated silencing of *imb-1* (left pair of boxes) or *imb-3* (right pair of boxes). (D) Total number of behavioural differences detected between *imb-2*(*syb6272*) and N2 reared from the L1 stage at higher temperatures (25°C, 27°C or 28°C, left 3 bars) or exposed to higher temperatures (28°C, 30°C or 32°C) for 4h prior to tracking (right 3 bars). Red dashed line denotes the number of phenotypic changes detected between *imb-2*(*syb6372*) and wild-type reared at 20°C without any treatment. Speed of worms upon stimulation with three 10 s pulses of high intensity blue light (blue shaded regions), each 100 s apart. Both strains were treated with 1 mM paraquat for 4h prior to tracking. Coloured lines represent average speed of the detected worms across all biological replicates and shaded areas represent 95% confidence intervals. (F-G) Key behavioural differences of strains treated with 1 mM paraquat (as described in 5E). (H) Fraction of worms moving after 4h exposure to 0.5 or 1 mM aldicarb. Individual points marked on the box plots are well averaged values (3 – 5 worms per well) for each feature across the independent days of tracking. *P*-values are for comparison to wild-type N2, treated with the same stressor condition, using block permutation t-tests (*n* = 10,000 permutations, correcting for multiple comparisons using the Benjamini-Yekutieli method).

Next, we used temperature as a candidate sensitiser. Rearing both strains from the L1 stage at higher temperatures (25°C, 27°C or 28°C, Figure 5D) leads to a reduction in the number of behavioural differences between the strains. Exposing adult worms to 28°C for 4h prior to tracking causes negligible difference in the separation of mutant and wild-type strains. 4h exposure of *imb-2[D157N]* and N2 to 32°C causes all worms to become near-stationary and there is little detectable difference. Although a comparison of worms exposed to 30°C for 4h enhances the *imb-2*(*syb6372*) phenotype, this may simply reflect a greater variability in these conditions since most of the differences relate to feature interquartile ranges rather than their medians.

Following temperature, we tested how chemical perturbation with paraquat or aldicarb affected behaviour. Paraquat is a herbicide known to induce oxidative stress and behavioural changes in *C. elegans*^83,84^. We find that exposure of mutant and wild-type strains to paraquat (1mM for 4h) results in 1907 significant behavioural differences between *imb-2*(*syb6372*) and N2. The majority of these new behavioural differences related to speed-associated features, with the mutant exhibiting a higher pre-stimulation (baseline) speed and a more sustained increase in speed following stimulation with blue light (Figure 5E). Furthermore, we see an increase in the overall duration of movement upon the blue light stimulation of *imb-2*(*syb6372*) (Figure 6F) and a greater separation of curvature related features (Figure 5G).

**Figure 6.**
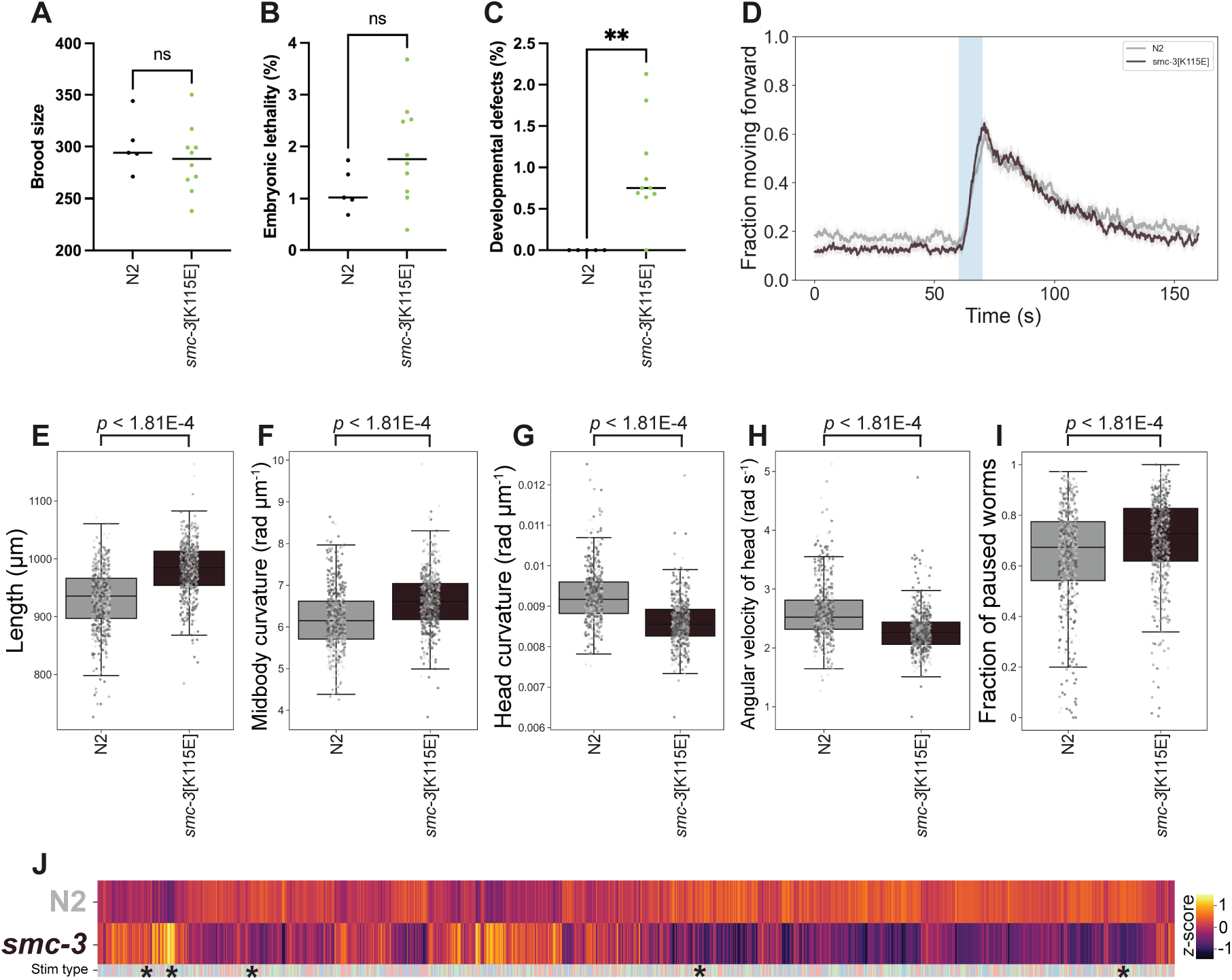
Patient-specific *smc-3[K115E]* mutants display behavioural defects. Quantification of brood size laid by wild-type controls (*n* = 5) and *smc-3 K115E* mutant worms (*n* = 10). *P* = 0.49 by a two-tailed Mann Whitney test. (B) Quantification of embryonic lethality amongst the broods displayed in (A). *P* = 0.10 by a two-tailed Mann Whitney test. (C) Quantification of the frequency of developmental defects amongst the broods displayed in (A). *P* = 0.0033 by a two-tailed Mann Whitney test. (D) Overall fraction of worms moving forward 60 s prior to and 80 s following stimulation with a 10 s blue light pulse (blue shading). Coloured lines represent averages of the detected fraction of paused worms across all biological replicates and shaded areas represent 95% confidence intervals. (E-I) Key behavioural features altered between N2 and the *smc-3 K11E* mutant. Individual points marked on the box plots are well averaged values (3 – 5 worms per well) for each feature across the independent days of tracking. *P*-values are for comparison to wild-type N2 using block permutation t-tests (*n* = 100,000 permutations, correcting for multiple comparisons using the Benjamini-Yekutieli method). (J) Heatmap of the entire set of 8289 behavioural features extracted by Tierpsy. The ‘stim type’ barcode denotes when during image acquisition a feature was extracted: pre-stimulation (pink), blue light stimulation (blue) and post-stimulation (green). Asterisks show the location of the selected features presented in E-I.

Treatment with aldicarb causes an accumulation of acetylcholine in the synaptic cleft of the neuromuscular junction, leading to eventual paralysis, and is commonly used to identify altered synaptic transmission in *C. elegans* mutants^85^. There is no difference in the baseline movement of solvent treated worms, yet there is a dose-dependent decrease in the fraction of worms moving (increased paralysis) upon aldicarb treatment (Figure 5H). Treatment with 1mM aldicarb for 4h causes near-complete paralysis of N2 while *imb-2[D157N]* mutants are more likely to move showing some aldicarb resistance. This suggests decreased levels of synaptic transmission for *imb-2[D157N]* variants as acetylcholine accumulates more slowly in the synaptic cleft and that aldicarb may be a useful sensitiser for phenotypic screens.

### Mutation in cohesin subunit SMC-3

Cornelia de Lange syndrome (CdLS) is a genetic syndrome characterised by variable neurodevelopmental defects, facial dysmorphism, upper limb anomalies and atypical growth^68^. Most CdLS patients carry heterozygous mutations in different subunits of the cohesin complex, a molecular motor that is involved in chromosome segregation, DNA repair, and 3D genome organization^69^. How mutations in cohesin subunits cause CdLS is not fully understood, but it is thought that reduced cohesin function caused by heterozygous mutations induces defects in 3D genome organisation that alter the expression of developmental genes^70^. In contrast, homozygous mutations that fully eliminate cohesin function are embryonically lethal, likely due to the essential role of cohesin in ensuring accurate chromosome segregation during cell division.

Genetic testing in individuals with clinical features that may be consistent with CdLS may identify *de novo* non-synonymous variants in cohesin subunits, including SMC3^71^. However, many identified variants are currently classified to be of uncertain clinical significance (VUS), for example if they have not been previously reported in other affected individuals. Often, the information that is currently available about the functional impact of a specific variant is limited.

To help explore this issue, we tested whether an SMC3[K114E] mutation identified as a variant of uncertain significance in an individual with a clinical presentation compatible with CdLS would cause any phenotypic defects in worms. The K114 residue is close to the ATPase head domain of SMC3 and corresponds to K115 in *C. elegans* SMC-3, thus we used CRISPR to create a *C. elegans* strain carrying the SMC-3[K115E] mutation. Similar to mammals, complete loss of cohesin function is lethal in *C. elegans* due to impaired chromosome segregation during embryonic development^72^. We first set up to determine whether the SMC-3 K115E mutations may represent a LoF mutation by investigating the brood size and viability of homozygous *smc-3[K115E]* mutants. Quantification of these two parameters revealed no significant differences between homozygous *smc-3[K115E]* mutant worms and wild-type controls (Figure 6A-B), suggesting that the SMC-3[K115E] mutation does not induce defects in chromosome segregation that compromise embryonic viability.

Despite this, we observed that a small, but significant, percentage of progeny from homozygous *smc-3[K115]* homozygous mutants displayed different developmental defects including: larval arrest, vulva protrusion, and abnormal body shape (Figure 6C). This suggested that the SMC-3[K115E] substitution does impact normal development, prompting us to investigate its potential effect on behavioural phenotypes. Tracking of adult homozygous *smc-3[K115]* mutant worms revealed strong behavioural phenotypes with 5685 significant behavioural differences compared to wild-type controls (Figure 6D-J). *smc-3[K115E]* worms are longer and have increased midbody curvature, but decreased curvature and agular velocity of the head. The patient avatar is less active during baseline recordings but has a normal response to stimulation with bluelight. These results demonstrate that the SMC-3[K115E] substitution in *C. elegans* causes developmental and behavioural defects without impacting on embryonic viability, mimicking the phenotype observed in CdLS patients and suggesting that the K114E in human SMC3 may contribute to the CdLS-like phenotypes observed in the heterozygous patient.

## Discussion

There are more than 3000 conserved *C. elegans* genes that currently have an association with human disease^7^. This number will grow as more patients are sequenced and the number of potential disease models is greater still since there may be multiple variants per gene that warrant separate modelling efforts. The high-throughput phenotyping of worm disease models provides an opportunity to map this extensive genotype-phenotype space.

Although we detected differences in all 25 of the strains in this study, three mutants (*imb-2*(*syb6372*), *shl-1*(*syb5907*), and *tmem-222*(*syb4882*)) showed weak to moderate phenotypes (<1000 significant feature differences) compared to wild-type animals. Strains with weak phenotypes are challenging to use in drug repurposing screens where the number of replicates per treatment is likely to be small. In previous work, a chemical sensitiser caused an easily measurable phenotype that was used in a successful drug repurposing screen^86^. We treated *imb-2(syb6372*) mutants with different temperatures, RNAi against related genes, and two compounds with different modes of action. We chose these methods because they are compatible with high-throughput screening. Other candidates for sensitisation include optogenetic stimulation of specific neural circuits and mechanical stimuli delivered to the entire plate. Other kinds of behavioural assays such as chemotaxis that might draw out novel phenotypes are more difficult to perform with high-throughput, although it is an area of active research^87^. RNAi resistance is a previously reported *imb-2* phenotype^88^, so it is perhaps unsurprising that we were unsuccessful in using targeted genetic sensitisation methods to enhance our *imb-2[D157N]* mutant phenotype. Nonetheless, we remain optimistic that the specificity of RNAi and the ease of delivery through feeding will be useful in other cases to sensitise disease models.

Likewise, chemical perturbation is easily incorporated into this phenotyping approach and may help to identify additional cellular pathways that are affected by disease mutations. For example, 4h aldicarb treatment led to a clear separation between *imb-2*(*syb6372*) and N2. This could enable future drug screens to be performed by identifying compounds that rescue the paralysis phenotype, hence pushing the behaviour of the mutant towards the healthy control. A caveat of this approach is that drugs that themselves caused paralysis or had broad toxic effects would show up as hits and need to be filtered with a counter screen. At the same time, aldicarb resistance in *imb-2[D157N]* worms suggests a slower accumulation of acetylcholine in the synaptic cleft^85^, a phenotype that would not have been obvious from tracking in unperturbed conditions. Similarly, we identify that FLCN deficiency may lead to an increased susceptibility of mTORC2 inhibition, whereas FNIP1/2 deficiency may cause a greater susceptibility to MTOR1/2 inhibition.

Overall, our results further support the use of high-throughput tracking to phenotype diverse worm disease models, extending our previous results on knockouts to more models, including patient-specific single amino acid changes and heterozygous mutants. Similar to the knockout strains, several of the point mutants showed strong phenotypes that could support drug repurposing screens. The relative ease and low cost of generating new worm models, coupled with the simplicity and ease of phenotyping them, means a systematic search for treatments to hundreds of Mendelian diseases is within reach.

## Materials and Methods

### Mutant generation

To generate loss-of-function mutants, CRISPR guide RNAs were designed to target large deletions that start close to the start codon and excise several exons from a gene to yield high confidence loss of function of the entire gene. For patient-specific allelic variants, guide RNAs were designed to target a single *C. elegans* codon corresponding to the position of the desired amino acid change in humans. Mutants were designed, made and confirmed by sequencing in N2 background by SunyBiotech. Transgenic worms carrying the patient variant SMC3 Lys114Glu were generated by in vitro assembly of Cas9 ribonucleoprotein complexes and a ssDNA repair template containing the desired amino acid change franked with short sequences (35 bp) homologous to the target region^89^. The ATATTACATTGATAATAAAA sgRNA was used to target the desired *smc-3* region.

### C. elegans preparation

For standard phenotyping experiments, all strains were cultured on Nematode Growth Medium (NGM) at 20°C and fed with *E. coli* OP50 following standard procedure^90^. Synchronised populations of young adult worms were used for imaging and were cultured by bleaching unsynchronised gravid adults and allowing L1 diapause progeny to develop for 2.5 days at 20°C (detailed protocol: https://dx.doi.org/10.17504/protocols.io.2bzgap6). *let-526*(*syb8759*) and *odr-8*(*syb4940*) were developmentally delayed and allowed to grow for longer (6h and 7h, respectively) before imaging. Before imaging, young adults were washed in M9 (detailed protocol: https://dx.doi.org/10.17504/protocols.io.bfqbjmsn) and transferred to tracking plates (3-5 worms per well) using a COPAS 500 Flow Pilot (detailed protocol: https://dx.doi.org/10.17504/protocols.io.bfc9jiz6), before being returned to a 20°C incubator for 3.5h. Plates were transferred onto the multi-camera tracker for another 30 minutes to habituate prior to imaging (detailed protocol: https://dx.doi.org/10.17504/protocols.io.bsicncaw).

### Compound screening

For the chemical treatment of *imb-2*(*syb6372*), aldicarb and paraquat were dissolved in ddH_2_O. The day prior to tracking, imaging plates were dosed to achieve the desired final well concentration of each compound (1 mM paraquat and 0.5 or 1 mM aldicarb) prior to seeding with bacteria (see below for details). Plates were left to dry (∼30min) and then stored in the dark at room temperature overnight. Using the methods above, *imb-2*(*syb6372*) and N2 worms (age synchronized young adults) were dispensed into the imaging plate wells and incubated at 20°C for 4h before tracking.

*E. coli* OP50 treated with 0.5% paraformaldehyde for 2h^64^, then washed 3 times in M9 buffer, was used as the food source for the folliculin mutant drug screens (detailed protocol:https://dx.doi.org/10.17504/protocols.io.81wgbyq71vpk/v1). All compounds acting on the AMPK/mTOR pathways were dissolved in DMSO. The day prior to tracking, imaging plates were dosed with the compounds to achieve a final well concentration of 100 μM prior to seeding with PFA-treated *E. coli* OP50. N2, *flcn-1*(*syb8071*) and *fnip-2*(*syb8038*) were all exposed to the compounds, or an identical volume of DMSO only (1% w/v), for 4h prior to tracking (as described above).

### Increased temperature screening

When comparing the behaviour of *imb-2*(*syb6372*) and N2 at increased temperatures, worm populations were age synchronised as described above. For rearing the strains at higher temperatures, L1 worms were refed onto plates seeded with *E. coli* OP50 and incubated at 25°C, 27°C or 28°C. Worms were then washed and dispensed into tracking plates and incubated for 3.5h at their rearing temperature. 30 minutes prior to imaging, plates were transferred to the multi-camera tracker and allowed to habituate. For the acute exposure of strains to increased temperatures, *imb-2*(*syb6372*) and N2 were reared at 20°C. On the day of tracking, young adults were dispensed into tracking plates that were incubated at 28°C, 30°C and 32°C for 4h before being immediately imaged.

### RNAi silencing of imb-1 and imb-3

*E. coli* HT115(DE3) expressing *imb-1* or *imb-3* target-gene dsRNA from the Ahringer RNAi Feeding Library^91^ were used as a food source on tracking and nursery plates used to rear age synchronised *imb-2*(*syb6372*) and N2 worms to adulthood. Both NGM and no-peptone NGM was supplemented with tetracycline (15 μg mL^-1^, Sigma) and ampicillin (100 μg mL^-1^, Sigma) for microbial selection, alongside 1 mM isopropylthio-β-galactoside (IPTG, Thermo Fisher) to induce dsRNA production. Bacterial cultures were grown overnight (37°C, 200 rpm shaking) in LB (Miller) supplemented with tetracycline and ampicillin. Cultures were used to inoculate fresh LB (OD_600_ 1.0) supplemented with 1mM IPTG and incubated at 37°C until OD_600_ 1.0 was achieved (∼2h). Density-normalised cultures were cooled to 20°C and used in place of *E. coli* OP50, all other methods remain the same.

### Imaging plate preparation

No-peptone NGM was used for worm tracking plates (detailed protocol: https://dx.doi.org/10.17504/protocols.io.bvian4ae). Briefly, 20 g agar (Difco) and 3 g NaCl was dissolved in 975mL ddH_2_O and autoclaved. Once molten agar was cooled to 50-60°C post-autoclave salts (1 mL CaCl_2_ [1M]; 1 mL MgSO_4_ [1M]; 25 mL KPO_4_ [1M, pH 6.0]) and cholesterol (1 mL of 5 mg mL^-1^ stock) were added. Then 200 μL of agar was dispensed into each well of 96-square well plates (Whatman UNIPLATE: WHAT-77011651) using an Integra VIAFILL (detailed protocol: https://dx.doi.org/10.17504/protocols.io.bmxbk7in). Poured plates were stored in an air-tight container, agar side up at 4°C until required. One day before imaging, plates were dried in a cabinet to lose 3-5% of their weight by volume. An Integra VIAFILL dispenser was then used to seed the wells of each plate with 5 μL of bacterial food (OD_600_ 1.0).

### Image acquisition, processing and feature extraction

All videos were recorded and features extracted using methods we have previously described in detail^7^. Briefly, videos were acquired at 25 frames per second in a temperature-controlled room (20°C) using a shutter time of 25 ms and a resolution of 12.4 μm/pixel. Three sequential videos were recorded: a 5-minute pre-stimulus video; a 6-minute blue light recorded with three 10-second pulses of high intensity blue light 100-seconds apart (starting after 60 seconds); and a 5-minute post-stimulus recording. Videos were segmented and tracked using Tierpsy Tracker^9^ and a convolutional neural network classifier was used filter out non-worm objects^8^. We further filtered the data to only keep worm skeletons with a length of 700-1300 μm and width of 20-200 μm before using Tierpsy Tracker’s viewer to exclude wells with visible contamination, agar damage, compound precipitation, or excess liquid from downstream analysis. A previously-defined set of 3076 behavioural features^10^ were extracted for each well in the three videos (pre-stimulus, blue light and post-stimulus) and the feature values were averaged to give a single feature vector for each well (*n* = 1).

### Statistical analysis

Significant differences in the behavioural feature sets extracted from each model strain were compared to the N2 reference, for each of the 3 video recordings, using block permutation t-tests (https://github.com/Tierpsy/tierpsy-tools521/python/blob/master/tierpsytools/analysis/statistical_tests.py). Python (version 3.9.2) was used to perform the analysis, using *n* = 100,000 permutations that were randomly shuffled within the independent days of image acquisition (to account for day-to-day variation across day replicates). The resulting *p*-values were corrected for multiple comparisons using the Benjamini-Yekutieli method, with a false discovery rate of 5%^92^. The same methods were used to compare differences in the behaviour of *imb-2*(*syb6372*) and N2 treated with the same compounds, RNAi or the same change in temperature. For the folliculin mutant drug screens, statistical differences were calculated using the Kruskal-Wallis test and correcting for multiple comparisons using the Benjamini-Yekutieli method, between the DMSO-only (untreated) and compound-treated strains. Heatmaps, cluster maps and principal component analysis of the extracted feature sets for each strain compared to the N2 reference were calculated using Seaborn (0.11.1) packages^93^. All scripts used for statistical analysis and the generation of figures are available here: https://github.com/Tom-OBrien/High-throughput-behavioural-phenotyping-of-25-C.-elegans-disease-models.

## Data Availability

Datasets (extracted features, tracking data, and metadata) produced in this study are available on Zenodo: https://zenodo.org/records/13941711.

## Supporting information

Strain Specific Gene Cards

## Acknowledgements

This project has received funding from the European Research Council (ERC) under the European Union’s Horizon 2020 research and innovation programme (Grant agreement No. 714853) and was supported by the Medical Research Council through grants MC-A658-5TY30 and MC-A654-5QB10. The work on *imb-2* was funded by the TNPO2 Foundation and the work on *let-526* by the Foundation for ARID1B Research. We would like to thank Reza Maroofian for advice on disease genes to model.

## Supplementary Information

**Supplementratry Figure 1.**
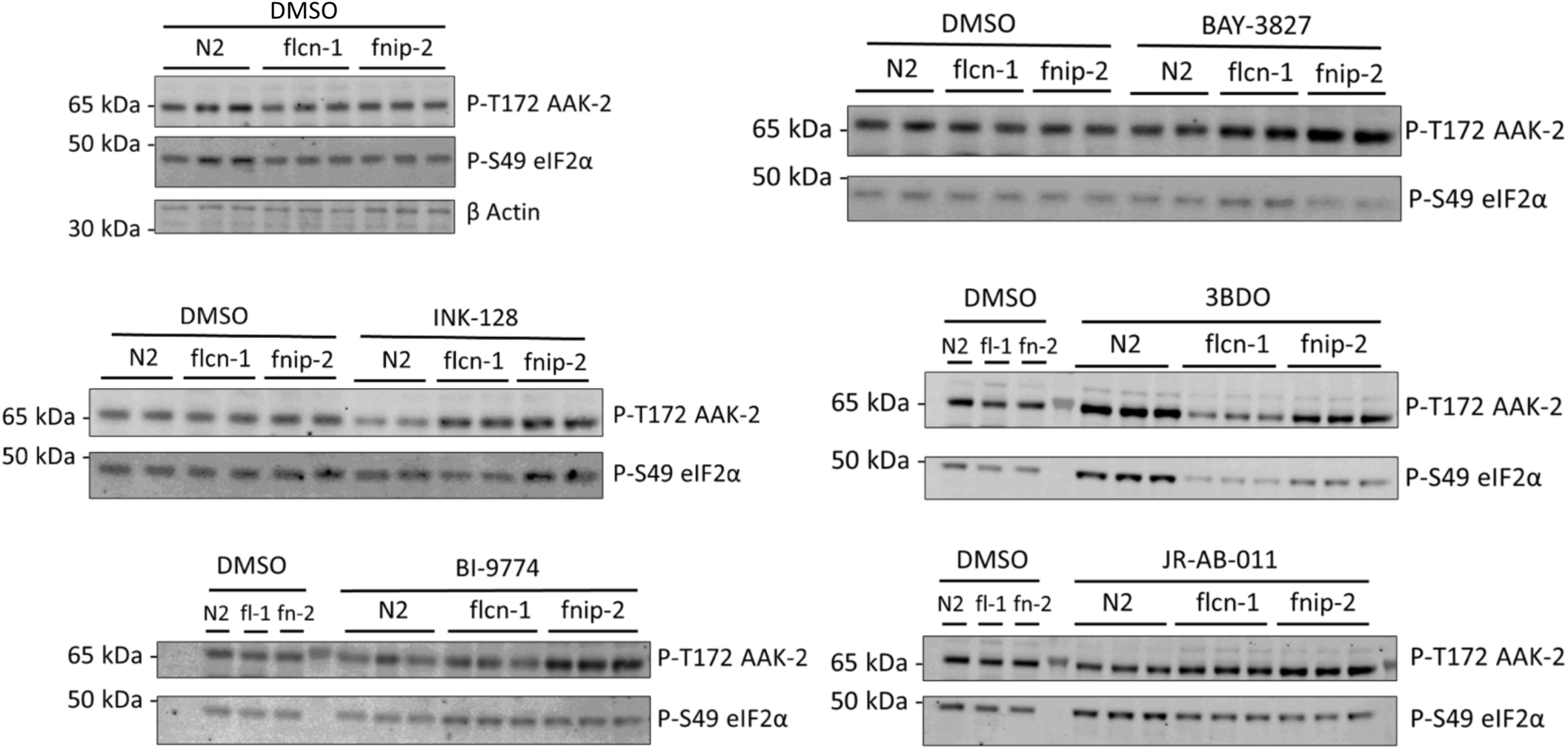
Western blots of mTOR and AMPK activity. Changes in AMPK and mTOR signalling were monitored using antibodies against P-T172 AMPK and P-S49 eIF2α respectively using protein lysates of N2, flcn-1 and fnip-2 worms treated with the indicated drugs at the same concentrations used for behavioural study.Blots show technical replicates representative of (*n* = 5), each condition was monitored with two blots, one shown here. Western blotting was performed as described in the supplementary methods. β Actin was used as loading control.

**Supplementary Figure 2.**
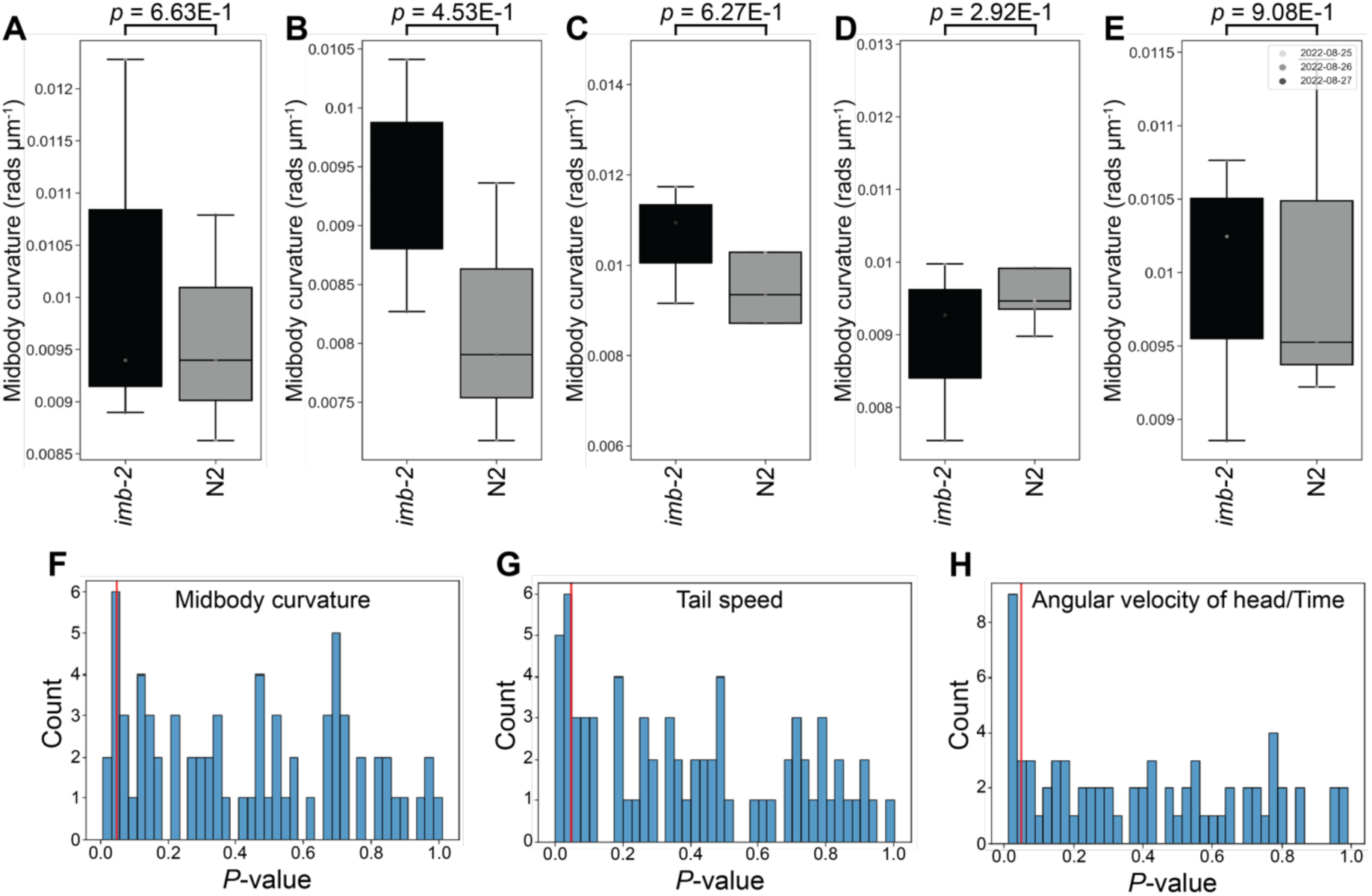
TNPO2 patient avatar phenotypes cannot be reliably detected with a low number of well replicates. (A-E) Sub-sampled boxplots of the same behavioural feature (midbody curvature) previously detected as being statistically significant between *imb-2*(*syb6372*) and wild-type when using a large number of replicates (*n* > 300, strain specific gene card). Individual plots show randomly sampled data points (*n* = 3) from the same complete dataset. *P*-values are for comparison of *imb-2*[D157N] mutant to N2 using Student’s t-test, correcting for multiple comparisons using the Benjamini-Yekutieli method. (F-H) Histograms of 3 key behavioural features (shown in the strain specific gene card) showing the calculated *P*-value of each feature when sub-sampling the overall dataset to achieve *n* = 3 replicates (using same methods as for the individual boxplots). Red line shows *p* = 0.05, and is considered statistically significant. When looking at a small number of samples (mimicking what may be typically collected for a low replicate screen across a large number of compounds), the behavioural phenotype of the disease model mutant cannot be reliably distinguished from wild-type.

**Supplementary Figure 3.**
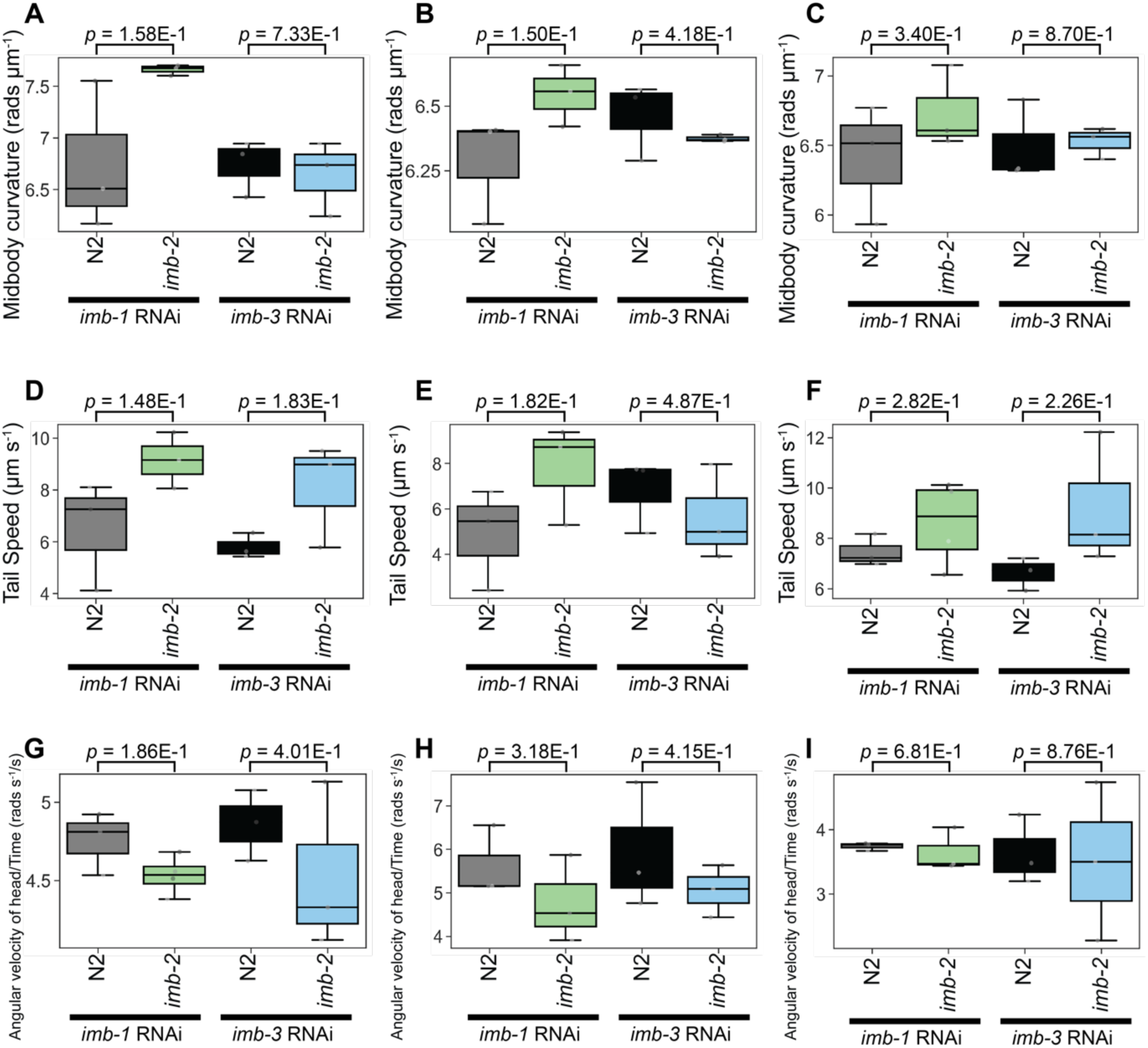
Targeted sensitisation of TNPO2 patient avatar does not cause a reliable phenotype when using a low number of well replicates. (A-I) Key behavioural phenotype of *imb-2*(*syb6372*) and wild-type worms following the RNAi-mediated silencing of *imb-1* (left pair of boxes) or *imb-3* (right pair of boxes). Individual plots show the same feature (repeated 3 times per row) for randomly sampled points (*n* = 3) from the complete dataset (shown in Figure 5). *P*-values are for comparison of *imb-2*[D157N] mutant to N2 using Student’s t-test, correcting for multiple comparisons using the Benjamini-Yekutieli method. As in Supplementary Figure 2a, when looking at a small number of samples the behavioural phenotype of the mutant cannot be reliably distinguished from wild-type.

## Supplementary Methods

### Antibodies

Phospho-AMPKα (Thr172) (40H9) Rabbit mAb #2535, Cell Signaling Technology

Phospho-eIF2α (Ser51) (119A11) Rabbit mAb #3597, Cell Signaling Technology

Beta Actin antibody 66009-1-Ig Clone 2D4H5, Proteintech

Goat anti-Mouse IgG Alexa Fluor Plus 800, Invitrogen A32730, Goat anti-Rabbit IgG Alexa Fluor Plus 680, Invitrogen A32734

### Buffers

#### MOPS running buffer

Invitrogen™ NuPAGE™ MOPS SDS Running Buffer (20X)

#### Tris-Glycine transfer buffer

25mM Tris, 192mM glycine

#### TBST

10mM Tris, pH8, 0.15M NaCl, 0.5% Tween20 (v/v)

#### 4x SLB

0.089 M Tris pH6.8, 13% (v/v) glycerol, 2.7% (v/v) SDS, 7% (v/v) beta mercaptoethanol, 1% (v/v) bromophenol blue)

#### 2X Lysis Buffer

100 mM Hepes pH7.4, 100 mM NaF, 10 mM NaPP, 2 mM EDTA, 20% (v/v) glycerol and 2% Triton X-100 (v/v), 2 mM DTT, 8 μg/ml trypsin inhibitor, 0.2 mM PMSF, Additionally, 1 PhosSTOP™ tablet and 1 cOmplete™, Mini, EDTA-free Protease Inhibitor Cocktail tablet per 10ml of buffer.

#### Protein extraction

Packed worm pellets were resuspended in 2x lysis buffer matching the volume of the pellet 1:1 Worm lysate mixes were transferred to 1.5ml tubes with metal beads and put in a bead mill homogenizer at 4 °C. Samples were homogenised at the highest speed for 4 min. Samples were spun down at 19,000 x g for 20 min at 4 °C to clarify the protein extract, this supernatant was used for western blotting.

#### Western Blotting

15μg of lysate supernatant per sample was used and boiled with 1x SLB for 3 minutes. SDS-PAGE was then carried out using a 4-12% Gradient Polyacrilamide gel Bis-Tris 1mm (Invitrogen NP0323BOX) in MOPS buffer at 180 volts constant voltage for 45 minutes, protein ladder used was PageRuler™ (ThermoScientific 26617). Protein transfer into a PVDF membrane was performed in Tris-Glycine transfer buffer at 4°C and 40V constant voltage overnight (16h), using a Bio-Rad PROTEAN® Trans-Blot® Module and using it as described in the manufacturer’s protocol for overnight transfer (4°C and 40V).

Membranes were blocked using 4% (w/v) Skim milk powder in Tris Buffered Saline Tween (TBST) for 1 hour then washed once in TBST for 5 minutes, TBST buffer was used to wash after each following step three times for 5 minutes each. Primary antibodies were all used at 1/1000 dilution, 4% BSA (w/v) in TBST. Membranes undergo 16 hours of incubation at 4°C with primary antibodies. After washing, fluorescently tagged secondary antibodies were used at 1/5000 dilution, 4% (w/v) BSA in TBST, membranes were incubated in them for 1 hour at room temperature. After washing, membranes were scanned using LICOR Odyssey CLx and fluorescence was measured at 680nm and 800nm.

