## Supplementary material for "High-throughput behavioural phenotyping of 25 *C. elegans* disease models including patient-specific mutations": Strain Specific Gene Cards

### *blos-1(syb6895)*

Strain name: PHX6895
Wild-type gene length: 945bp
Mutation: 927bp deletion starting at nt position 9 (deletion of entire gene coding region)

#### *C. elegans* Description

<u>Function</u>: Involved in endosomal transport and lysosomal biogeniesis[1]; predicted to be part of the BLOC-1 complex

<u>Expression</u>: Excretory cell; intestine and nervous system

<u>Previously reported phenotypes</u>: None

#### Human Orthologs:

*BLOC1S1*

<u>Associated disease(s):</u>
Hermansky-Pudlak syndrome

#### Results

3927/8289 significant features vs N2 ($p < 0.05$, block permutation t-test, 10000 permutations)

Key phenotype(s): Shorter; decreased angular velocity, curvature, speed and acceleration of head; increased forward, but attenuated backward, photophobic escape response

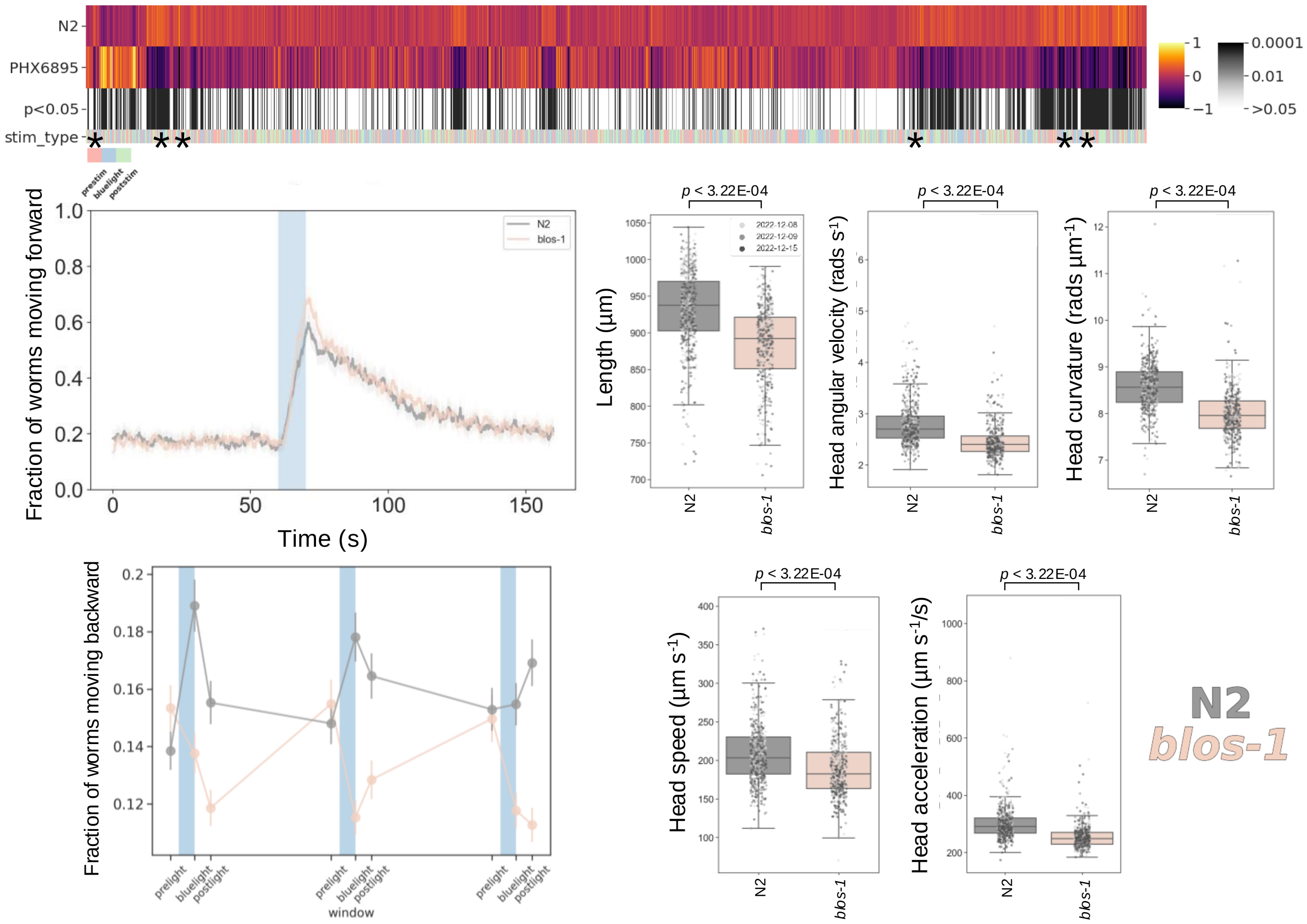

### *blos-8(syb6686)*

Strain name: PHX6686
Wild-type gene length: 1280bp
Mutation: 179bp deletion starting at nt position 10 (deletion of entire gene coding region)

#### *C. elegans* Description

<u>Function</u>: Predicted to be a member of the BORC complex; associated with the cytosolic face of lysosomes[2]

<u>Expression</u>: Unknown

<u>Previously reported phenotypes</u>: None

#### Human Orthologs:

*BORCS7*

<u>Associated disease(s):</u>
Hermansky-Pudlak syndrome; Hereditary spastic paraplegia

#### Results

3034/8289 significant features vs N2 ($p < 0.05$, block permutation t-test, 10000 permutations)

Key phenotype(s): Longer; decreased angular velocity of hips; increased midbody curvature; decreased tail instantaneous angular acceleration (time derivative of angular velocity); decreased sinusoidal undulations (time derivative of midbody curvature); attenuated backward photophobic escape response

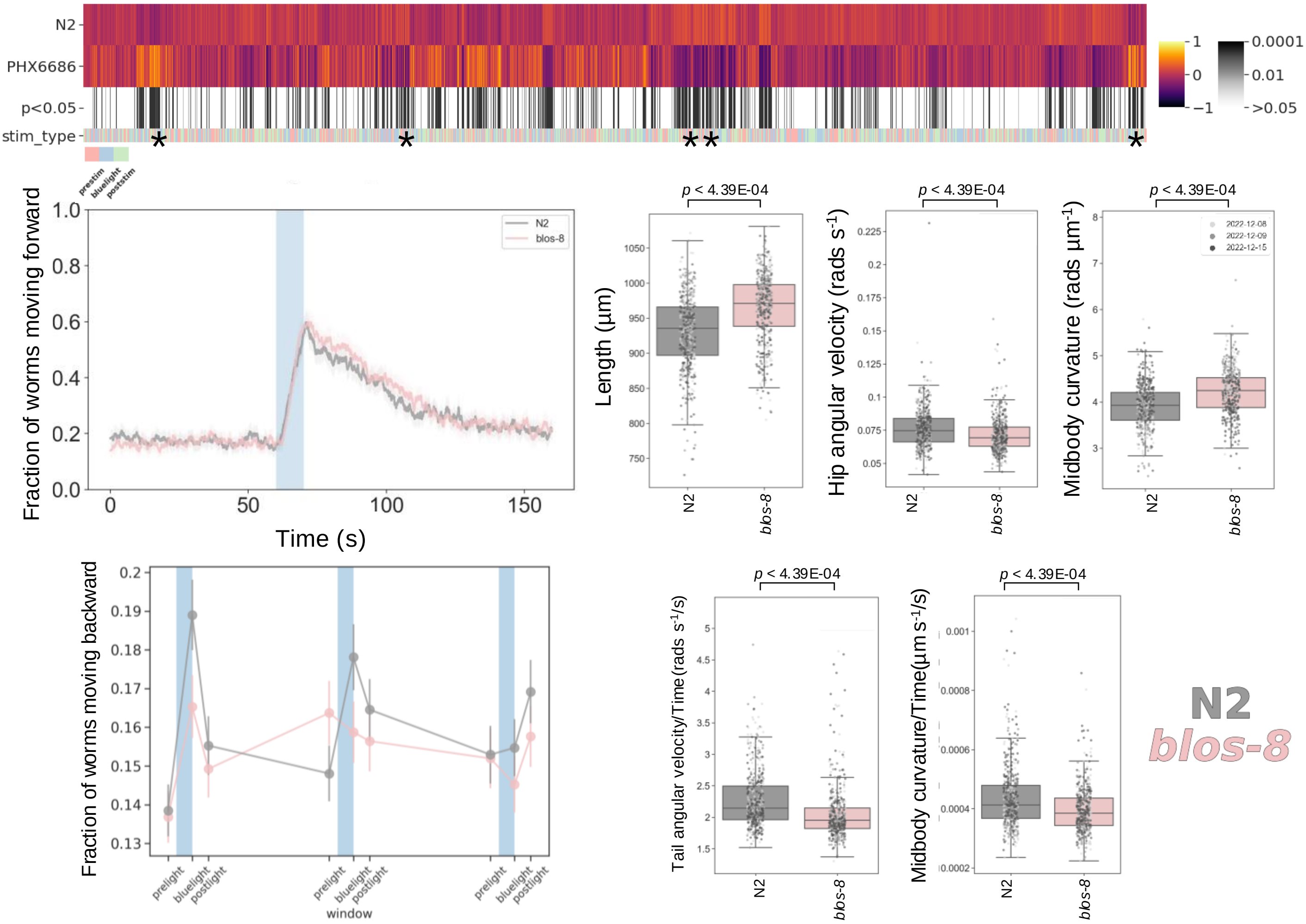

### *blos-9(syb7029)*

Strain name: PHX7029
Wild-type gene length: 1046bp
Mutation: 945bp deletion starting at nt position 49 (half of exon 1 and remaining coding regions)

#### *C. elegans* Description

Function: Predicted to be a member of the BORC complex; associated with the cytosolic face of lysosomes[2]

Expression: Unknown

Previously reported phenotypes: None

#### Human Orthologs:

*BORCS8*

Associated disease(s):
Hermansky-Pudlak syndrome; Infantile-onset neurodegenerative disorder with altered lysosome dynamics

#### Results

3924/8289 significant features vs N2 ($p < 0.05$, block permutation t-test, 10000 permutations)

Key phenotype(s): Shorter; decreased angular velocity, curvature, speed and acceleration of head; attenuated forward, but increased backward, photophobic escape response

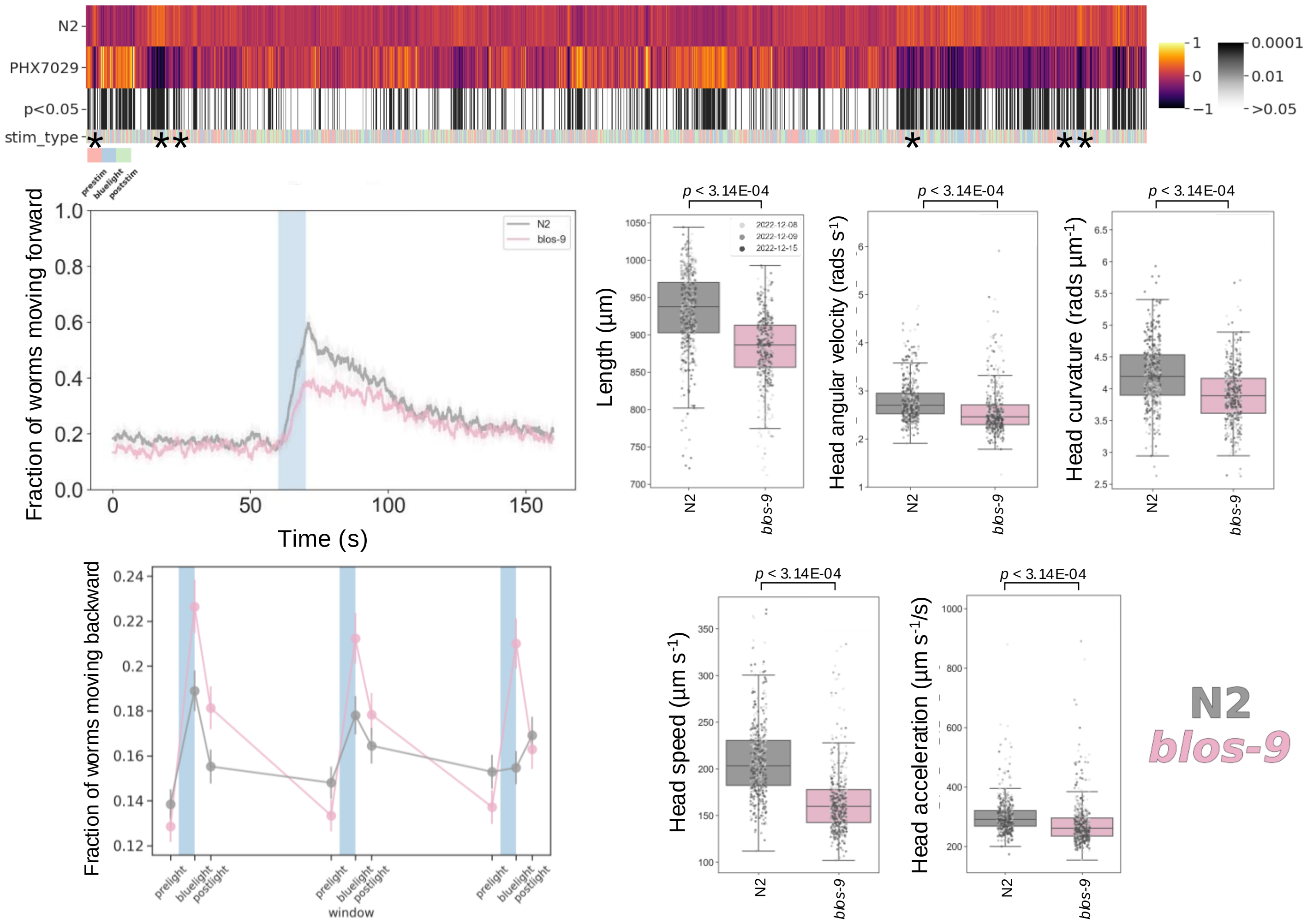

### *ccpp-1(syb4843)*

Strain name: PHX4843
Wild-type gene length: 6201bp
Mutation: 6180bp deletion starting at nt position 21 (deletion of entire gene coding region)

#### *C. elegans* Description

Function: Predicted to enable metallocarboxypeptidase activity and zinc ion binding activity; required for ciliary maintenance but not ciliogenesis[3]

Expression: Ciliated neurons and gubernacular muscle. Predicted to be located in cilium; dendrite and perikaryon

Previously reported phenotypes: Defective ciliary localization[4]; reduction of male mating efficiency and response of males to touch[4]

#### Human Orthologs:

*AGBL1*

Associated disease(s):
Fuchs' endothelial dystrophy

#### Results

2592/8289 significant features vs N2 ($p < 0.05$, block permutation t-test, 10000 permutations)

Key phenotype(s): Shorter; decreased angular velocity of midbody, neck and tail while moving backwards; lower frequency midbody bends (time derivative of midbody curvature); decreased overall speed; attenuated backing in response to blue light (but no change in forward photophobic escape response)

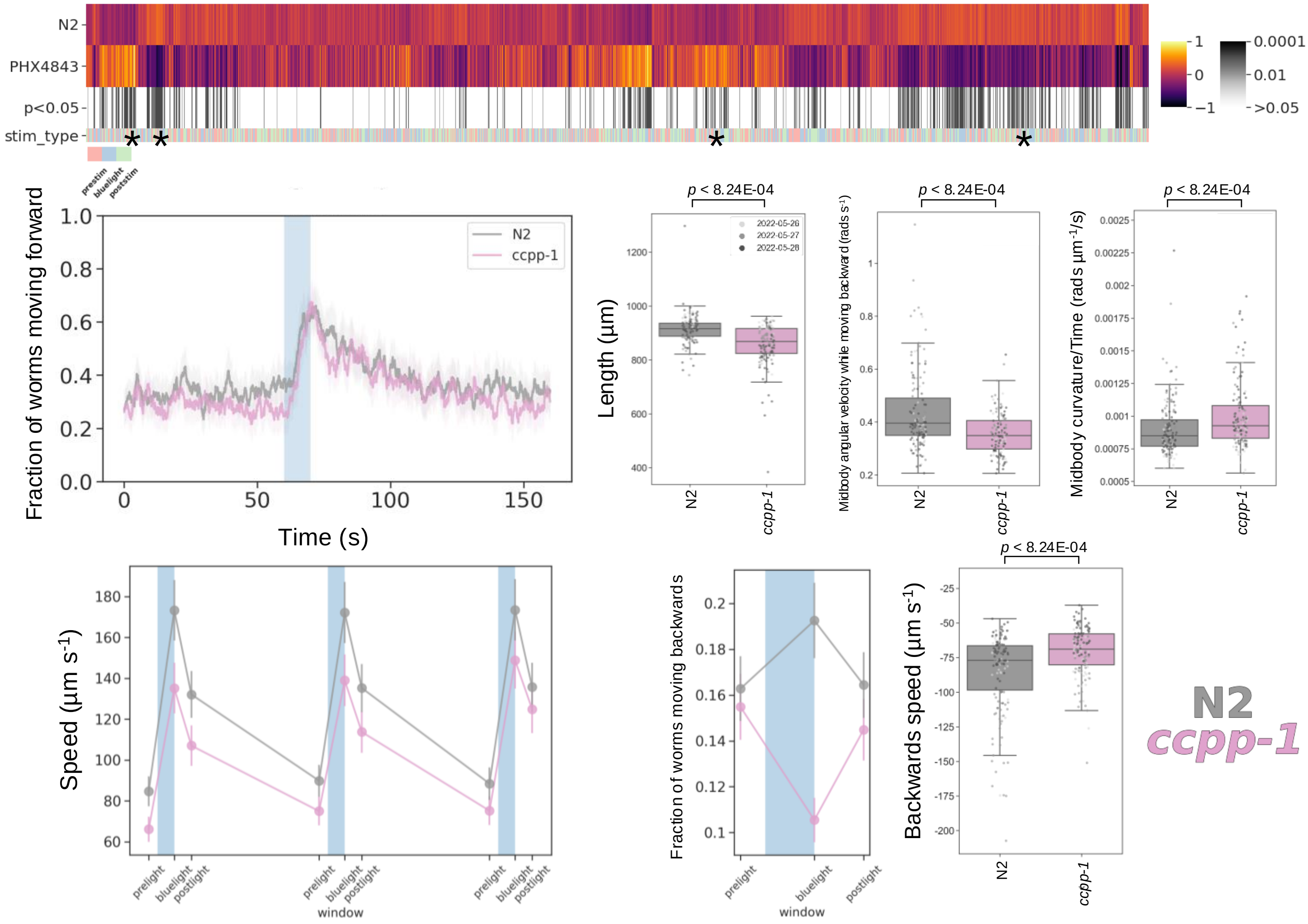

### *cpx-1(syb7694)*

Strain name: PHX7694
Wild-type gene length: 1568bp
Mutation: 1196bp deletion starting at nt position 41 (deletion of entire gene coding region)

#### *C. elegans* Description

Function: Enables SNARE binding activity; involved in the regulation of neurotransmitter secretion and synaptic vesicle exocytosis

Expression: Head neurons; motor neurons; tail neurons; ventral cord neurons

Previously reported phenotypes: aldicarb hypersensitivity[5]; defective for thrashing in liquid[5]

#### Human Orthologs:

*CPLX1 and CPLX2*

Associated disease(s):
Developmental and epileptic encephalopathy; infantile myoclonic epilepsy; intellectual disability; Alzheimer's disease; Parkinson's disease

#### Results

5801/8289 significant features vs N2 ($p < 0.05$, block permutation t-test, 10000 permutations)

Key phenotype(s): Shorter; less active, with attenuated photophobic escape response; increased angular velocity and decreased curvature (for all body segments); decreased speed; increased frequency of body bends (time derivative of midbody curvature)

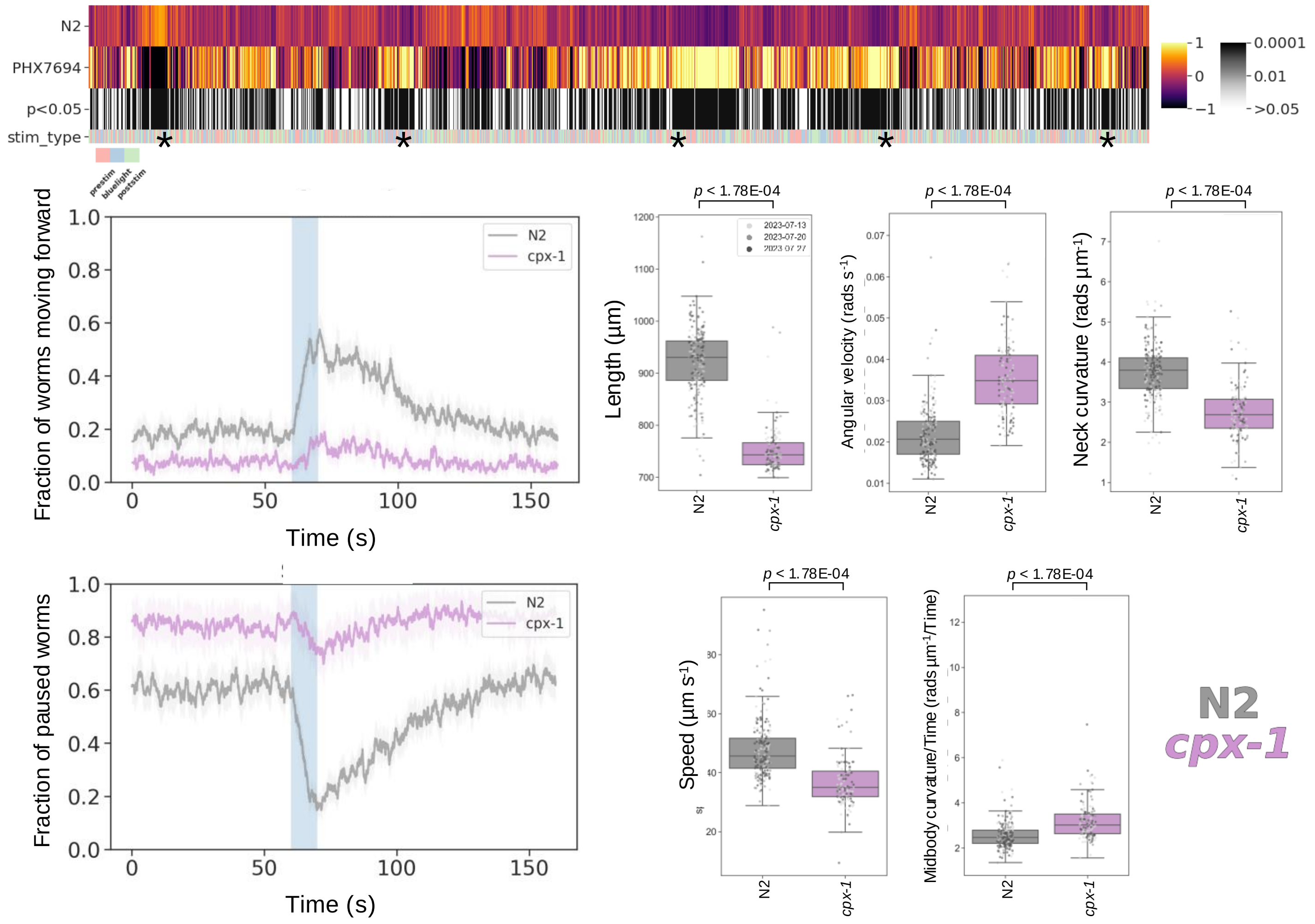

### *flcn-1(syb8071)*

Strain name: PHX8071
Wild-type gene length: 18087bp
Mutation: 17582bp deletion starting at nt position 296 (from start of exon 1 to end of exon 8)

#### *C. elegans* Description

Function: Contributes towards GTPase activator activity; negative regulation of RNA polymerase II transcription; positive regulation of TORC1 signalling and transforming growth factor beta receptor signalling

Expression: excretory cell; nervous system; spermatheca; vulva

Previously reported phenotypes: Increased resistance to hyperosmotic stress[6]; increased glycogen content[6]; increased lifespan[7]

#### Human Orthologs:

*FLCN*

Associated disease(s):
Birt-Hogg-Dube syndrome; primary spontaneous pneumothorax; renal cell carcinoma

#### Results

3663/8289 significant features vs N2 ($p < 0.05$, block permutation t-test, 10000 permutations)

Key phenotype(s): Hyperactive; increased duration of forward and backward locomotion in response blue light stimulation; increased speed; increased angular velocity (all body segments); deeper and more frequent body bends (time derivative of angular velocity or curvature, respectively)

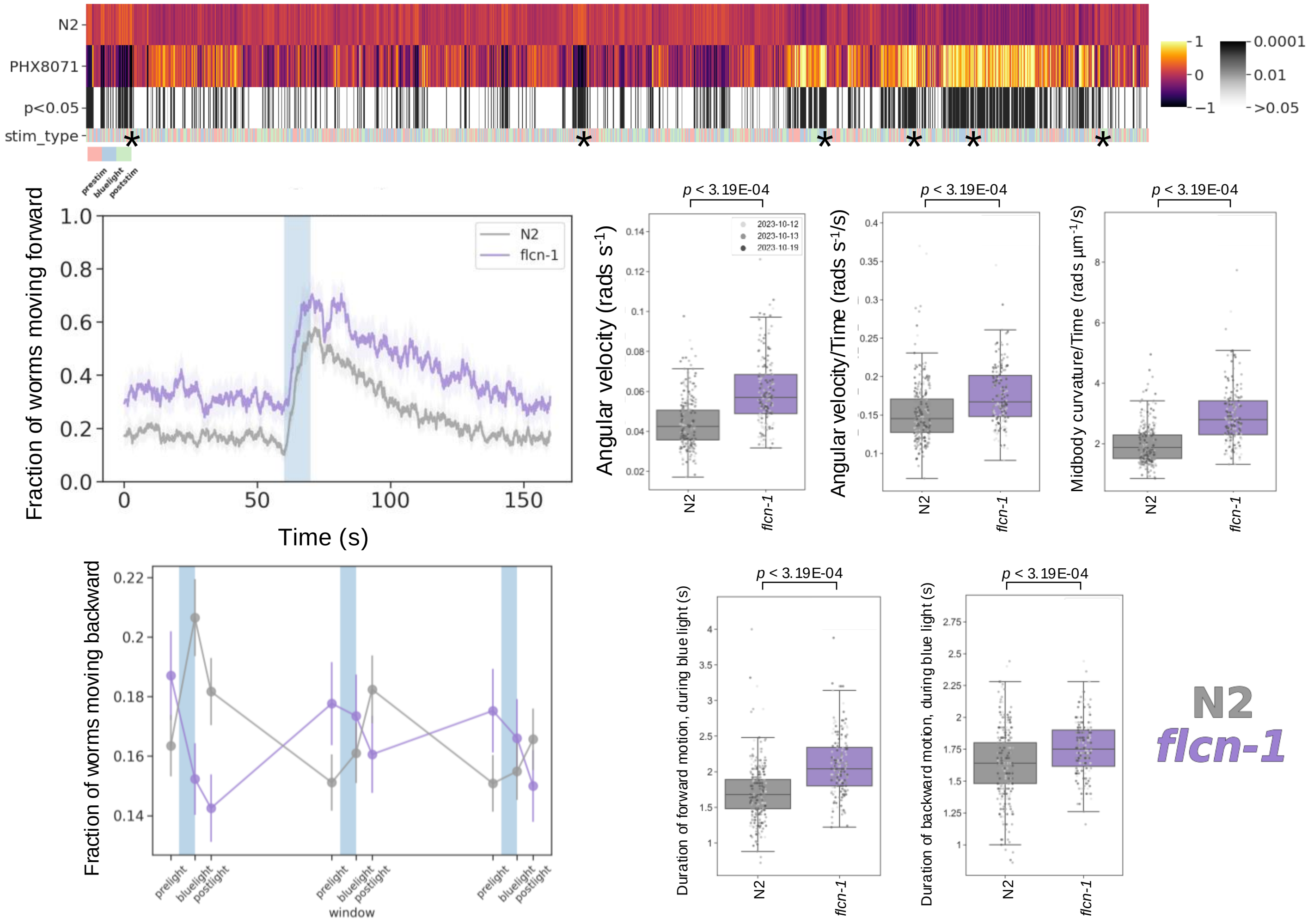

### *fnip-2(syb8038)*

Strain name: PHX8038
Wild-type gene length: 13091bp
Mutation: 12685bp deletion starting at nt position 135 (deletion of entire gene coding region)

#### *C. elegans* Description

<u>Function</u>: Enables ATPase inhibitor activity and protein-folding chaperone binding activity

<u>Expression</u>: Predicted to be located in cytoplasm

<u>Previously reported phenotypes</u>: None

#### Human Orthologs:

*FNIP1 and FNIP2*

<u>Associated disease(s):</u>
Primary immunodeficiency disease; obesity

#### Results

3433/8289 significant features vs N2 ($p < 0.05$, block permutation t-test, 10000 permutations)

Key phenotype(s): Hyperactive; short; decreased duration of backward locomotion in response blue light stimulation; increased angular velocity (all body segments); increased speed; deeper and more frequent body bends (time derivative of angular velocity or curvature, respectively)

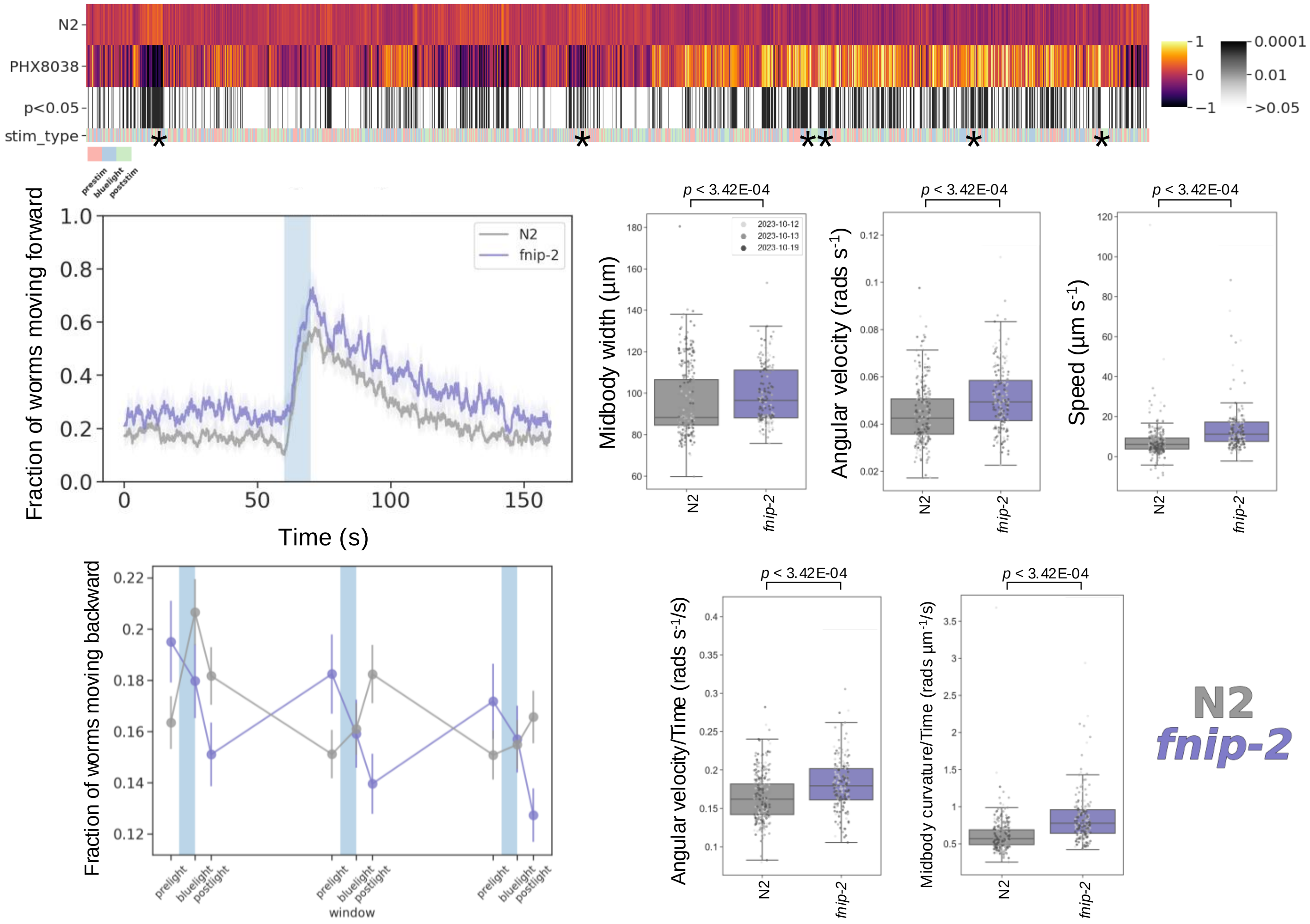

### ***imb-2(syb6372)***

Strain name: PHX6372 *imb-2*(*syb6372*)[D157N]
Wild-type gene length: 931bp
Mutation: D to N point mutation at amino acid position 157. Synonymous to D156N in TNPO2

#### *C. elegans* Description

Function: Nuclear import signal receptor activity; nuclear localisation sequence binding activity; protein import into nucleus

Expression: AWC$^{ON}$, AECL and AWCR sensory neurons

Previously reported phenotypes: Reduction of odour chemotaxis[8]

#### Human Orthologs:

*TNPO2*

Associated disease(s):
Intellectual developmental disorder with hypotonia; Impaired speech; Dysmorphic facies

#### Results

770/8289 significant features vs N2 ($p < 0.05$, block permutation t-test, 10000 permutations)

Key phenotype(s): No difference in morphology or locomotion; decreased angular velocity of head; increased curvature of midbody and tail; decreased head foraging (time derivative of angular velocity); decreased frequency of pausing (during stimulation with blue light); increased tail speed

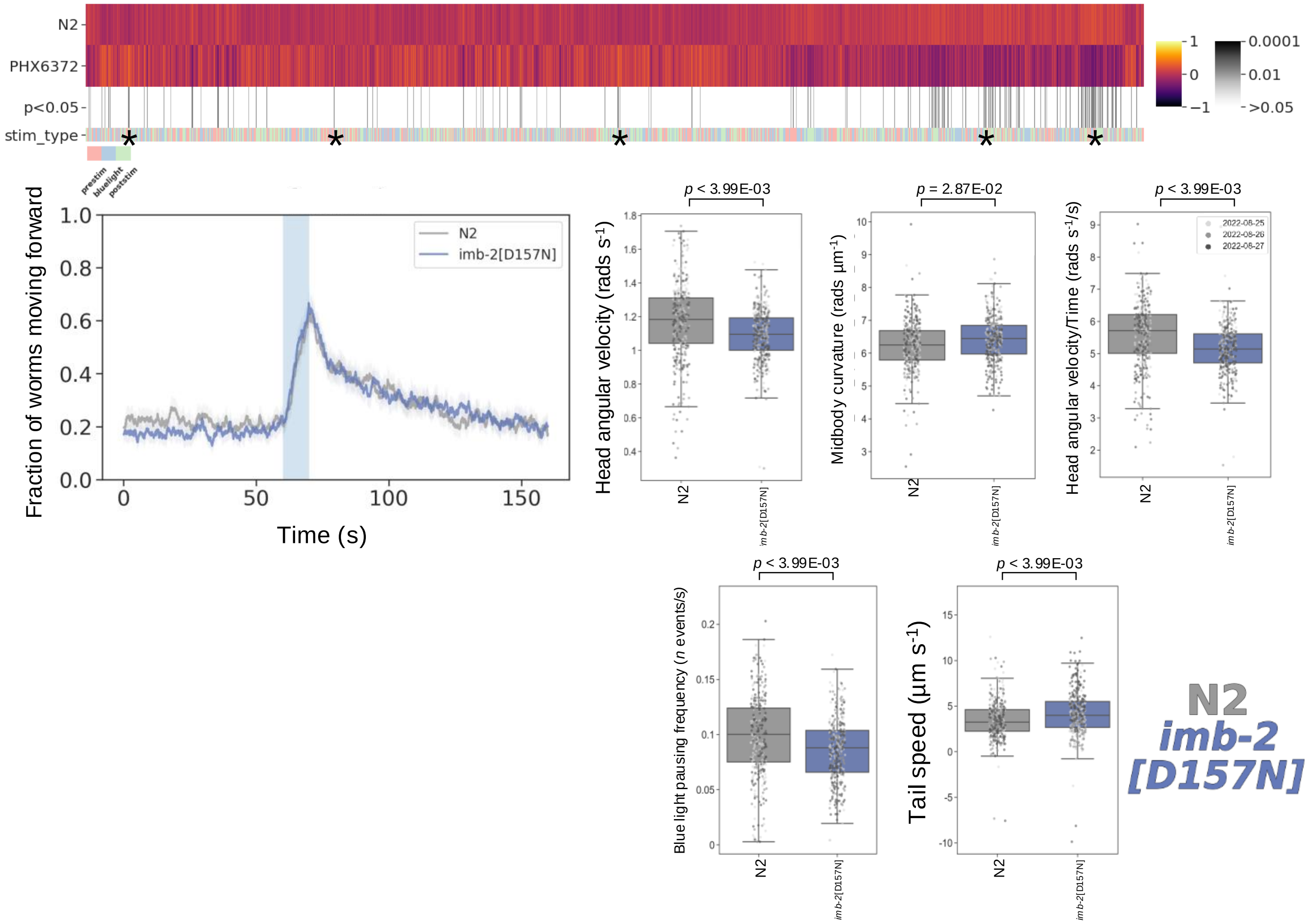

### *irk-1(syb5903)*

Strain name: PHX5903
Wild-type gene length: 11931bp
Mutation: 11338bp deletion starting at nt position 168 (starting at beginning of exon 1)

#### *C. elegans* Description

Function: Enables inward rectifier potassium channel activity[9]; regulation of G protein-coupled receptor signalling[9]; regulation of egg-laying behaviour[9]; inhibition of HSNs by EGL-6 signaling[9]

Expression: Egg-laying apparatus; gonad; intestine; and neurons

Previously reported phenotypes: Mild defect in egg laying[10]

#### Human Orthologs:

*KCJN2; KCNJ3 and KCNJ4*

Associated disease(s):
Glucose metabolism disease; Heart conduction disease; Long QT syndrome

#### Results

3612/8289 significant features vs N2 ($p < 0.05$, block permutation t-test, 10000 permutations)

Key phenotype(s): Fatter; less active (high fraction of paused worms); decreased angular velocity of head while moving; lower frequency of head bends (time derivative of curvature); decreased; decreased head speed and overall speed of mutant

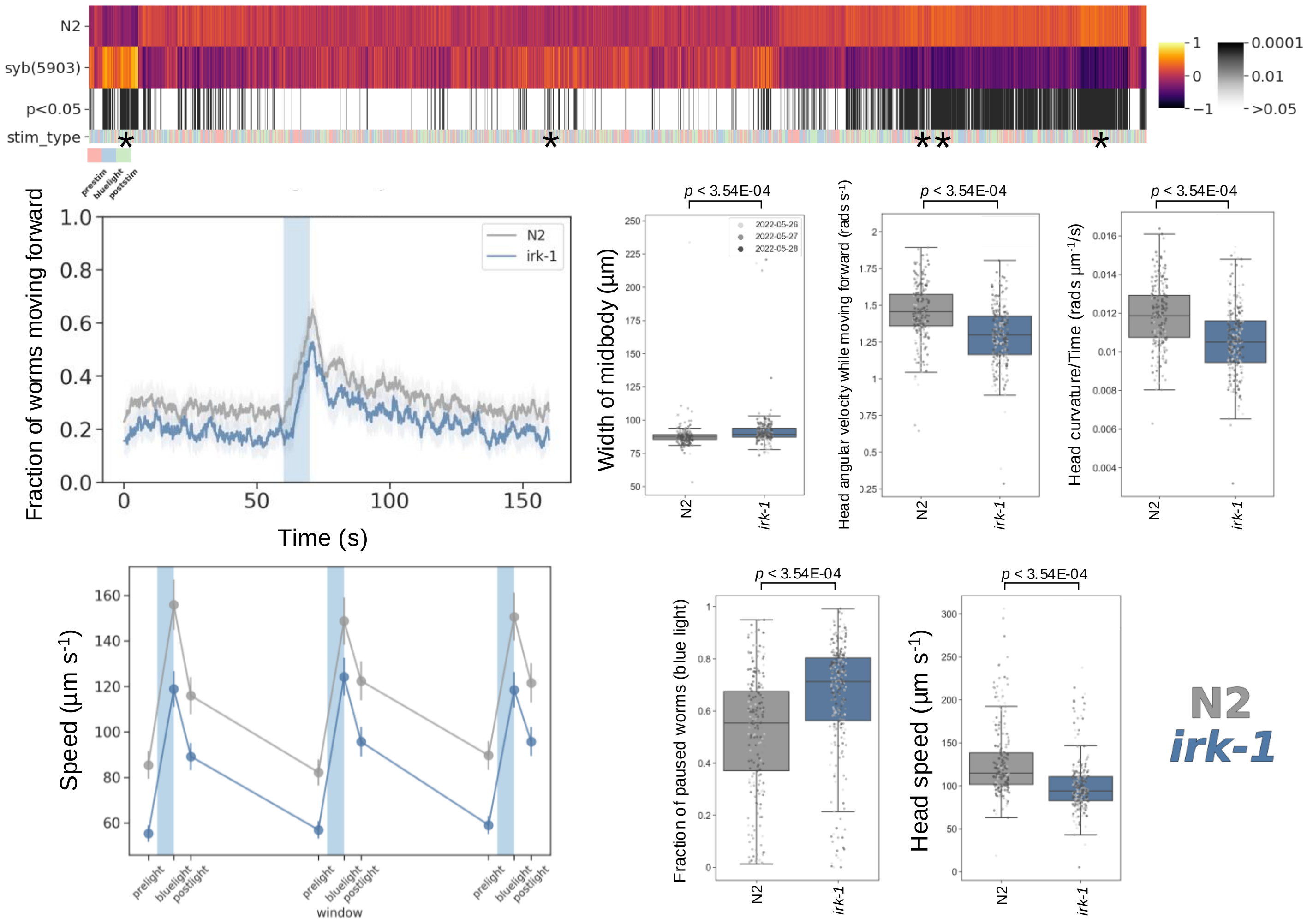

### *let-526(syb8759)*

Strain name: PHX8759 *let-526*(*syb8759*)I/hT2[*bli-4*(*e937*)*let-?*(*q782*)*qIs48*] (I;III)
Wild-type gene length: 1003bp
Mutation: Balanced heterozygote containing a 9024bp deletion starting at nt position 91 (deletion of entire gene coding region)

#### *C. elegans* Description

Function: Enables DNA binding activity; contributes to nucleosome binding activity; involved in gonad and larval development; positive regulation of transcription by RNA polymerase II

Expression: Excretory cell; gonad; pharynx; tail precursor cell; and vulva

Previously reported phenotypes: Sterility (homozygotes)[11] ;defective migration of excretory cell precursor[11]; PVD structural abnormalities[12]; perturbations in dendritic patterning and cell fate specification[12]

#### Human Orthologs:

*ARID1A* and *ARID1B*

Associated disease(s):
Coffin-Siris syndrome; Carcinoma (multiple); Pre-eclampsia

#### Results

5008/8289 significant features vs N2 ($p < 0.05$, block permutation t-test, 10000 permutations)

Key phenotype(s): 6 hour developmental delay; fatter; no difference in baseline locomotion, but attenuated photophobic escape response; decreased duration of movement in response to blue light; decreased angular velocity (all body segments); decreased curvature of head and neck; slower; lower frequency of head bends/foraging (time derivative of head curvature)

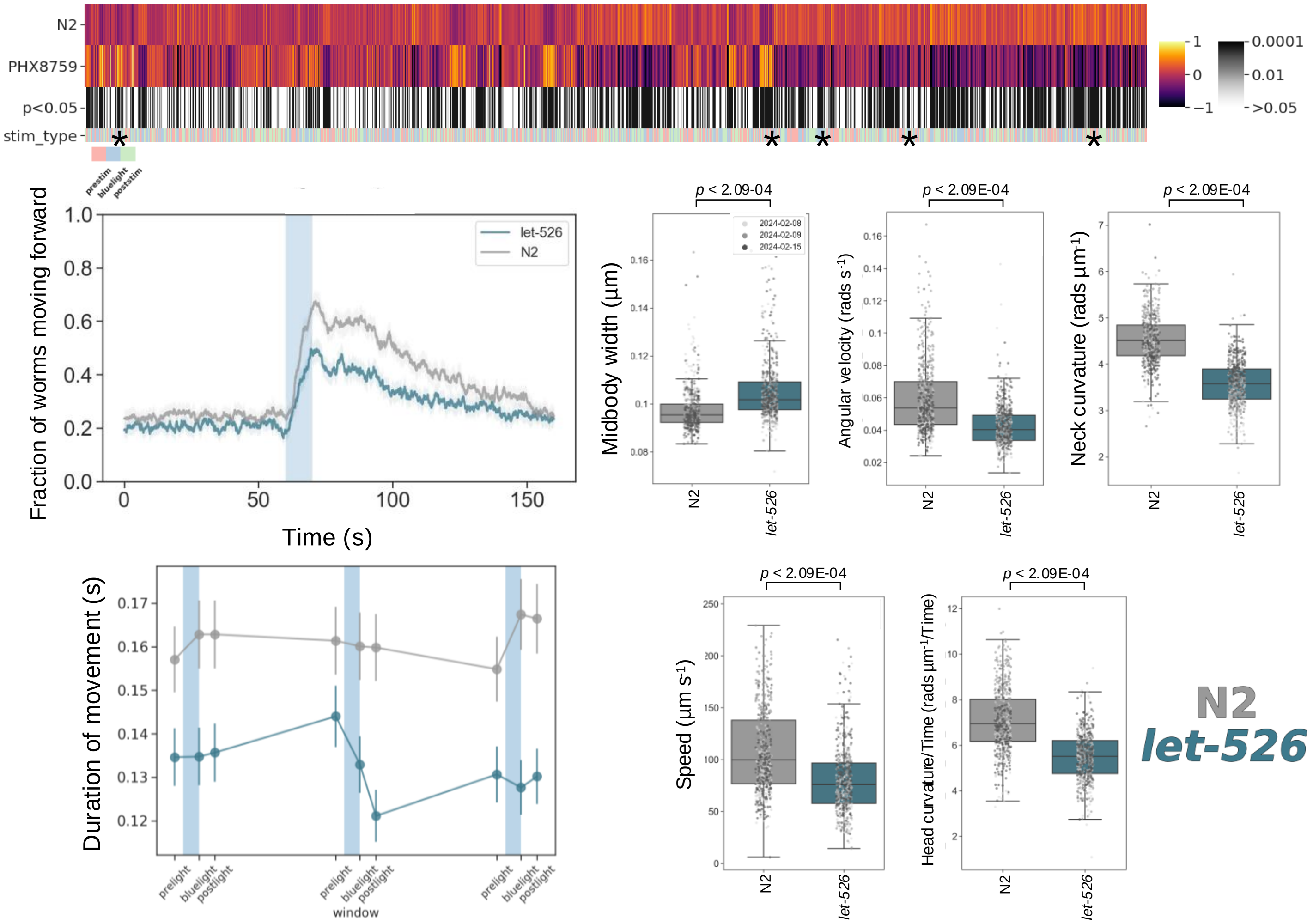

### *ncap-1(syb4935)*

Strain name: PHX4935
Wild-type gene length: 4624bp
Mutation: 4252bp deletion starting at nt position 166 (starting at first half of exon 1)

#### *C. elegans* Description

Function: Predicted to be involved in vesicle-mediated transport negative regulators of the AP2 clathrin adaptor complex[13]

Expression: Nerve ring

Previously reported phenotypes: None

#### Human Orthologs:

*NECAP1 and NECAP2*

Associated disease(s):
Developmental and epileptic encephalopathy (21)

#### Results

1826/8289 significant features vs N2 ($p < 0.05$, block permutation t-test, 10000 permutations)

Key phenotype(s): Shorter; decreased angular velocity, but increased curvature, of all body segments; less active; attenuated photophobic escape response; slower

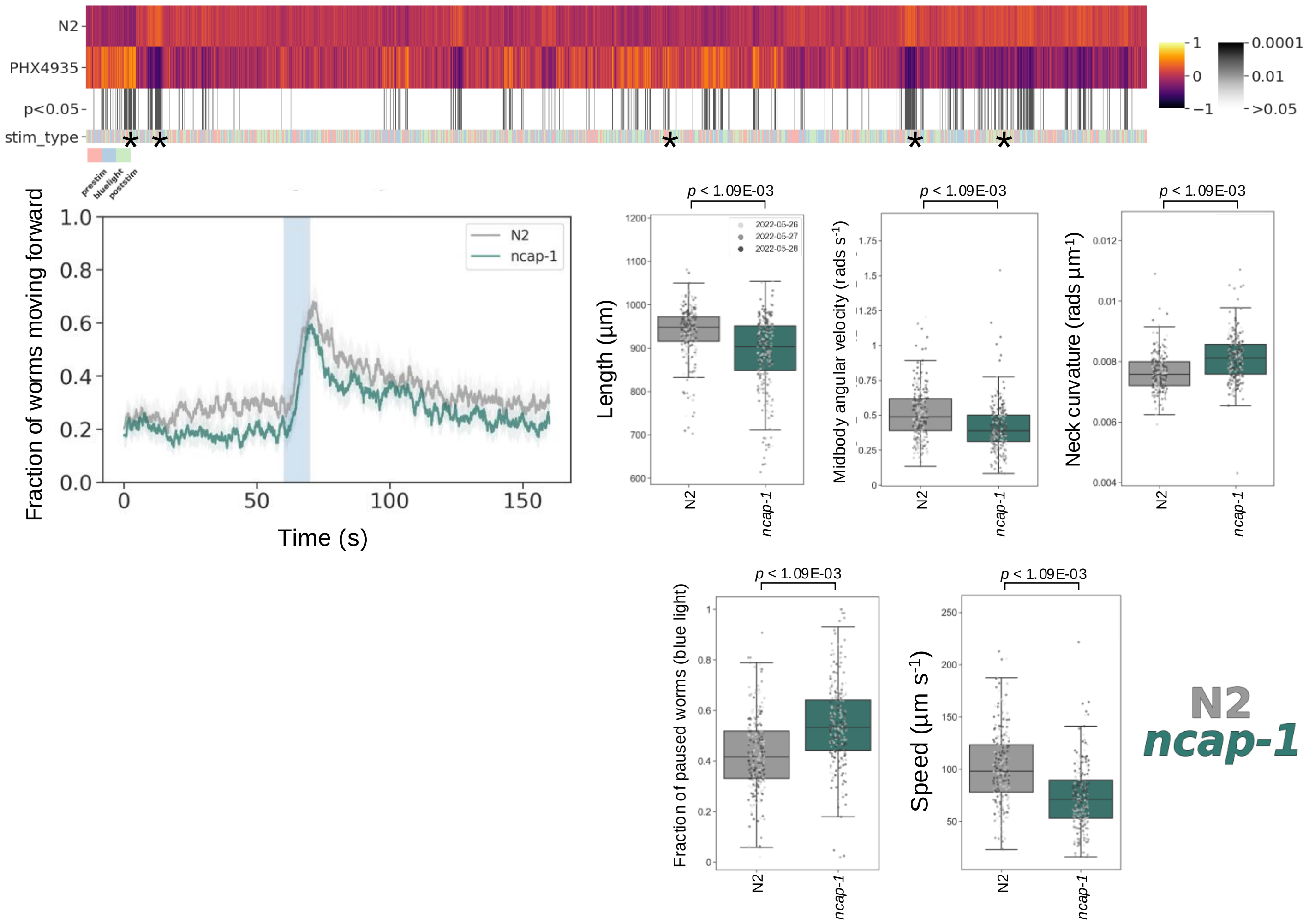

### *odr-8(syb4940)*

Strain name: PHX4940
Wild-type gene length: 3171bp
Mutation: 2585bp deletion starting at nt position 39 (starting from first half of exon 1)

#### *C. elegans* Description

Function: Enables deUFMylase activity; positive regulation of chemotaxis; protein localisation to non-motile cilium

Expression: Amphid and phasmid neurons

Previously reported phenotypes: AWA odorant chemotaxis defective[14]; diacetyl chemotaxis defective[15]; aldicarb hypersensitivity[15]

#### Human Orthologs:

*UFSP2*

Associated disease(s):
Early infantile epileptic encephalopathy; Developmental epileptic encephalopathy; Beukes hip dysplasia

#### Results

4557/8289 significant features vs N2 ($p < 0.05$, block permutation t-test, 10000 permutations)

Key phenotype(s): Developmental delay (7h); less active; shorter; decreased curvature (for all body segments); slower; decreased duration of forward movement, but a greater and more sustained backwards escape response to photophobic stimulation

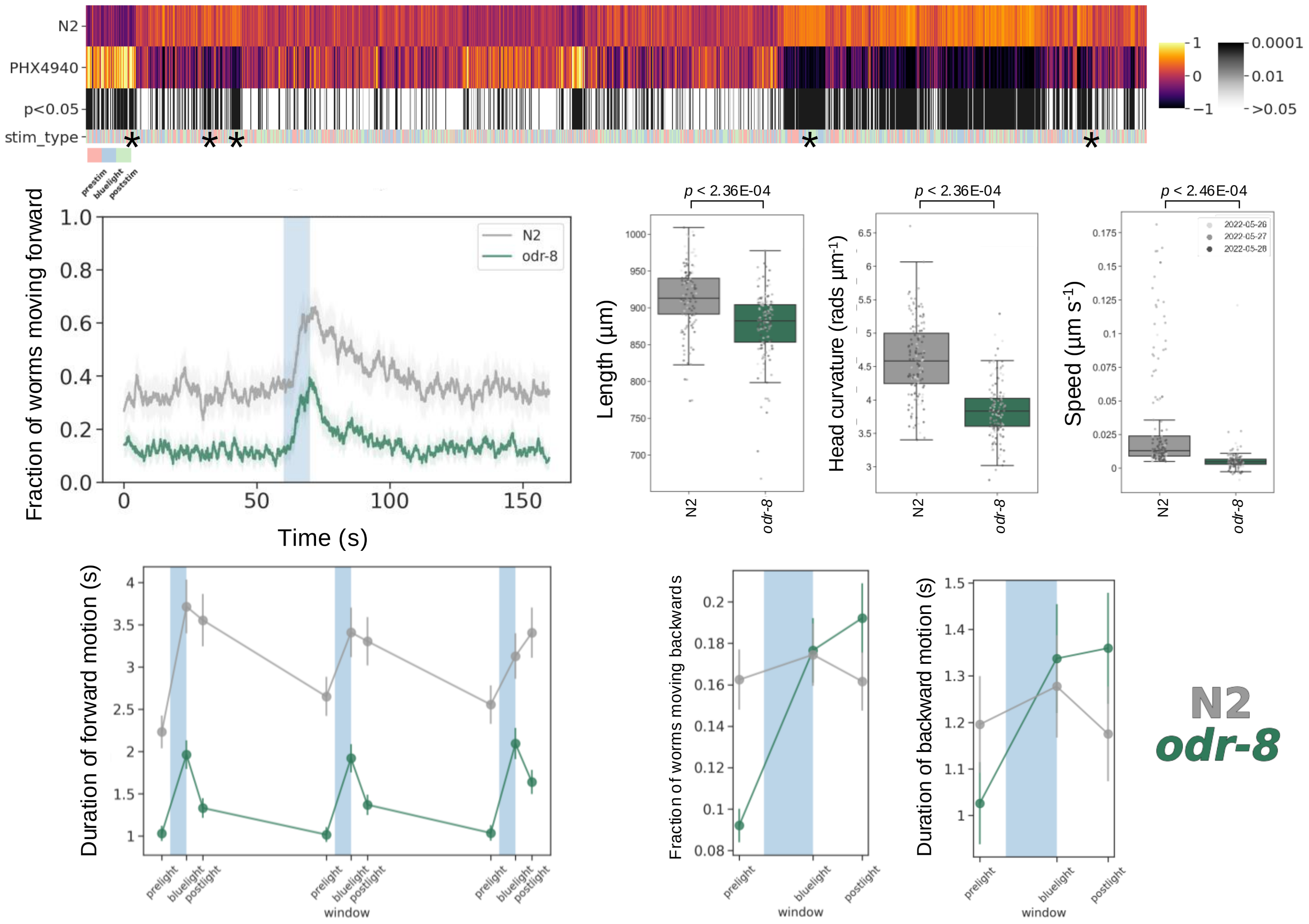

### *pacs-1(syb7634)*

Strain name: PHX7634 *pacs-1*(*syb7634*)[E205K]
Wild-type gene length: 1166bp
Mutation: E to K amino acid substitution at position 205- Synonymous to E209K mutation in human PACS2

#### *C. elegans* Description

Function: Involved in protein localisation to plasma membrane

Expression: Excretory cell; intestine; neurons; pharynx; vulva

Previously reported phenotypes: Aldicarb resistance[16]

#### Human Orthologs:

*PACS1 and PACS2*

Associated disease(s):
Schuurs-Hoeijmakers Syndrome; Developmental and epileptic encephalopathy (66); PACS1 syndrome; Seizures; Developmental delay; Intellectual disability; Autism

#### Results

3667/8289 significant features vs N2 ($p < 0.05$, block permutation t-test, 10000 permutations)

Key phenotype(s): Less active; decreased angular velocity; decreased curvature of head, neck, hips and midbody; lower frequency of body bends (time derivative of midbody curvature); decreased speed; attenuated photophobic escape response

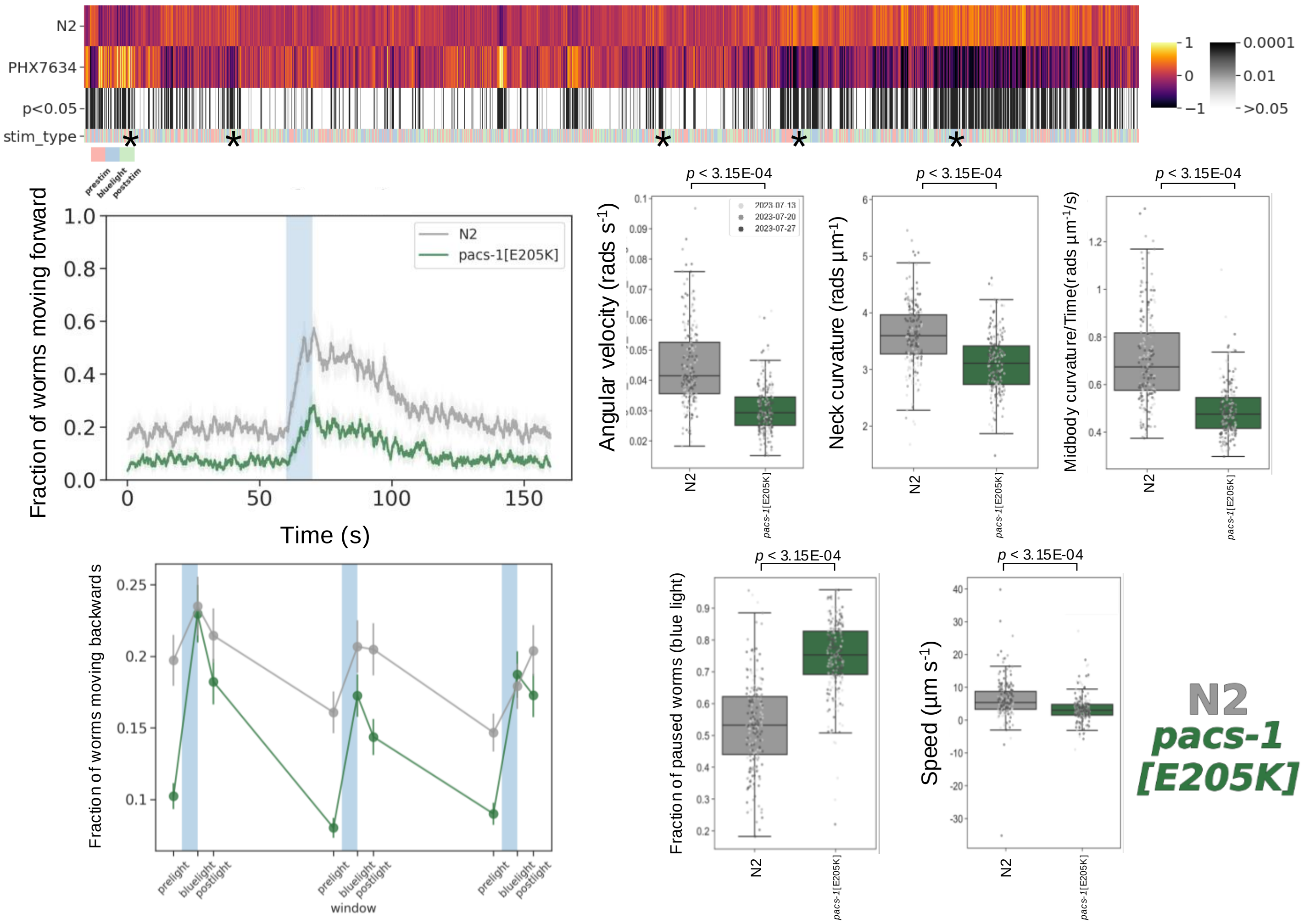

### *pde-1(syb7700)*

Strain name: PHX7700
Wild-type gene length: 7830bp
Mutation: 7095bp deletion starting at nt position 40 (deletion of entire gene coding region)

#### *C. elegans* Description

Function: Enables 3',5'-cyclic-AMP phosphodiesterase, 3',5'-cyclic-GMP phosphodiesterase and calmodulin binding activity; negative regulation of intracellular signal transduction; determination of adult lifespan; response to alkaline pH

Expression: Head; AFDL and AFDR neurons

Previously reported phenotypes: Isothermal tracking behaviour variant[17]; odorant adaptation variant[18]

#### Human Orthologs:

*PDE1A* and *PDE1B*

Associated disease(s):
Autosomal dominant nonsyndromic deafness (74); Cardiac disorders (multiple); Impotence; Schizophrenia

#### Results

3502/8289 significant features vs N2 ($p$ < 0.05, block permutation t-test, 10000 permutations)

Key phenotype(s): Shorter; fatter; decreased curvature (all body segments); lower frequency of body bends (time derivative of midbody curvature); less active during baseline recordings; shorter duration of movement in response to blue light; no difference in baseline speed, but decreased speed during photophobic escape response

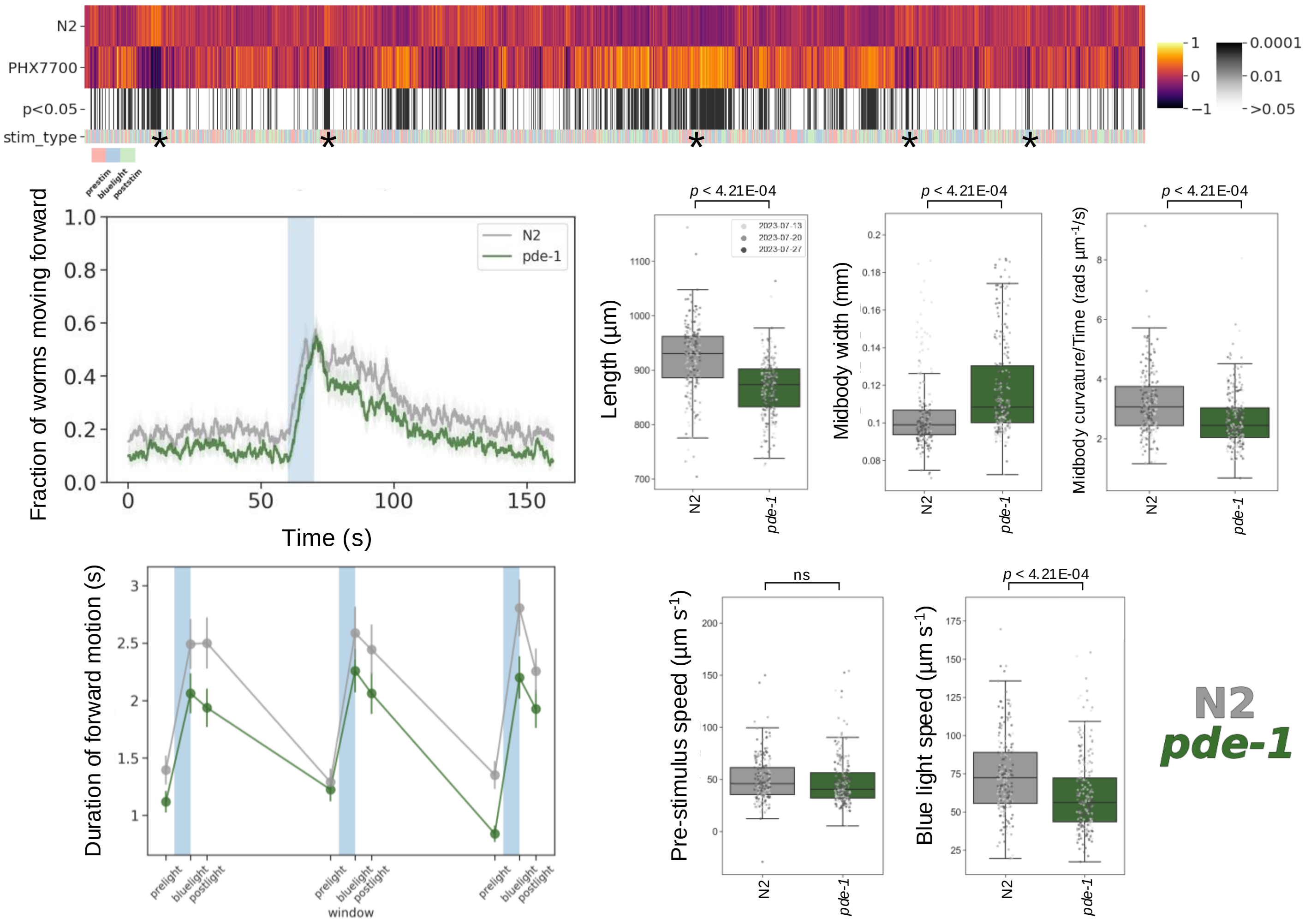

### *pde-5(syb7506)*

Strain name: PHX7506
Wild-type gene length: 4025bp
Mutation: 4025bp deletion of the entire gene

#### *C. elegans* Description

<u>Function</u>: Predicted to enable 3',5'-cyclic-GMP phosphodiesterase activity and cGMP-stimulated cyclic-nucleotide phosphodiesterase activity; negative regulation of cGMP-mediated signalling ;determination of adult lifespan; response to alkaline pH

<u>Expression</u>: AFDL and AFDR neurons

<u>Previously reported phenotypes</u>: Isothermal tracking behaviour variant[17]; odorant adaptation variant[18]

#### Human Orthologs:

*PDE10A*

<u>Associated disease(s):</u>
Schizophrenia; Huntington's disease; Dyskinesia; Striatal degeneration; Addiction

#### Results

3492/8289 significant features vs N2 ($p < 0.05$, block permutation t-test, 10000 permutations)

Key phenotype(s): Longer and fatter; most behavioural phenotypes affect the head only (location of gene expression), including: decreased angular velocity, curvature and speed, with lower frequency of head bends (time derivative of curvature); general locomotion is unaffected

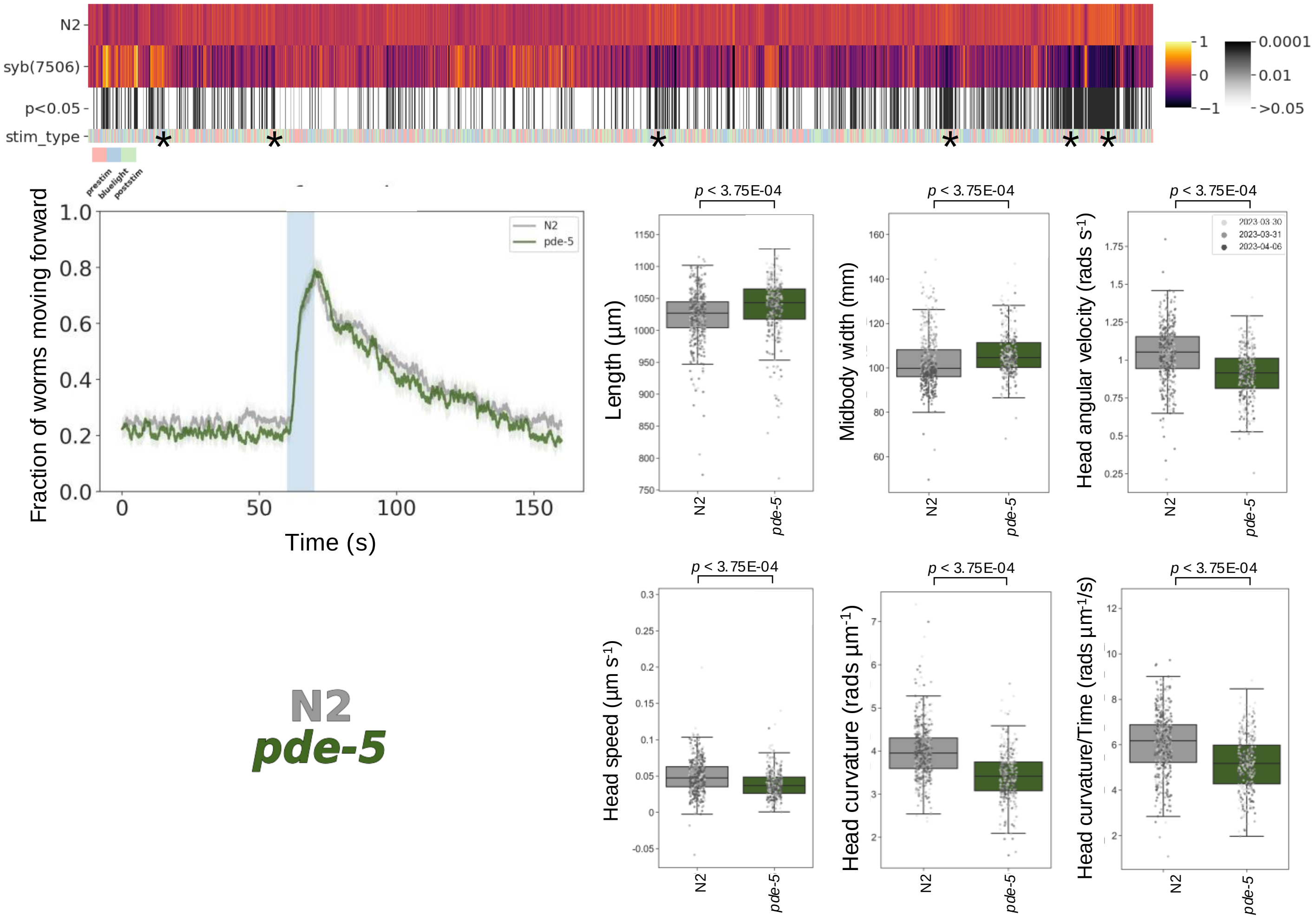

### ***pmp-4(syb7777)***

Strain name: PHX
Wild-type gene length: 3047bp
Mutation: 3047bp deletion of the entire gene

#### *C. elegans* Description

<u>Function</u>: Enables ATP binding activity; ATPase-coupled transmembrane and long-chain fatty acid transporter activity; involved in the fatty acid catabolic process, long-chain fatty acid import into peroxisomes and peroxisome organisation

<u>Expression</u>: Peroxisomal membrane (predicted)

<u>Previously reported phenotypes</u>: Decreased cilia length[19]; dye filling defect[19]; defective nonanone chemotaxis[19]

#### Human Orthologs:

*ABCD1* and *ABCD2*

<u>Associated disease(s):</u>
Adrenoleukodystrophy

#### Results

3329/8289 significant features vs N2 ($p < 0.05$, block permutation t-test, 10000 permutations)

Key phenotype(s): Shorter; increased angular velocity of tail and neck; lower frequency of tail bends (time derivative of tail angular velocity); decreased curvature of neck and midbody; no statistically significant changes in locomotion

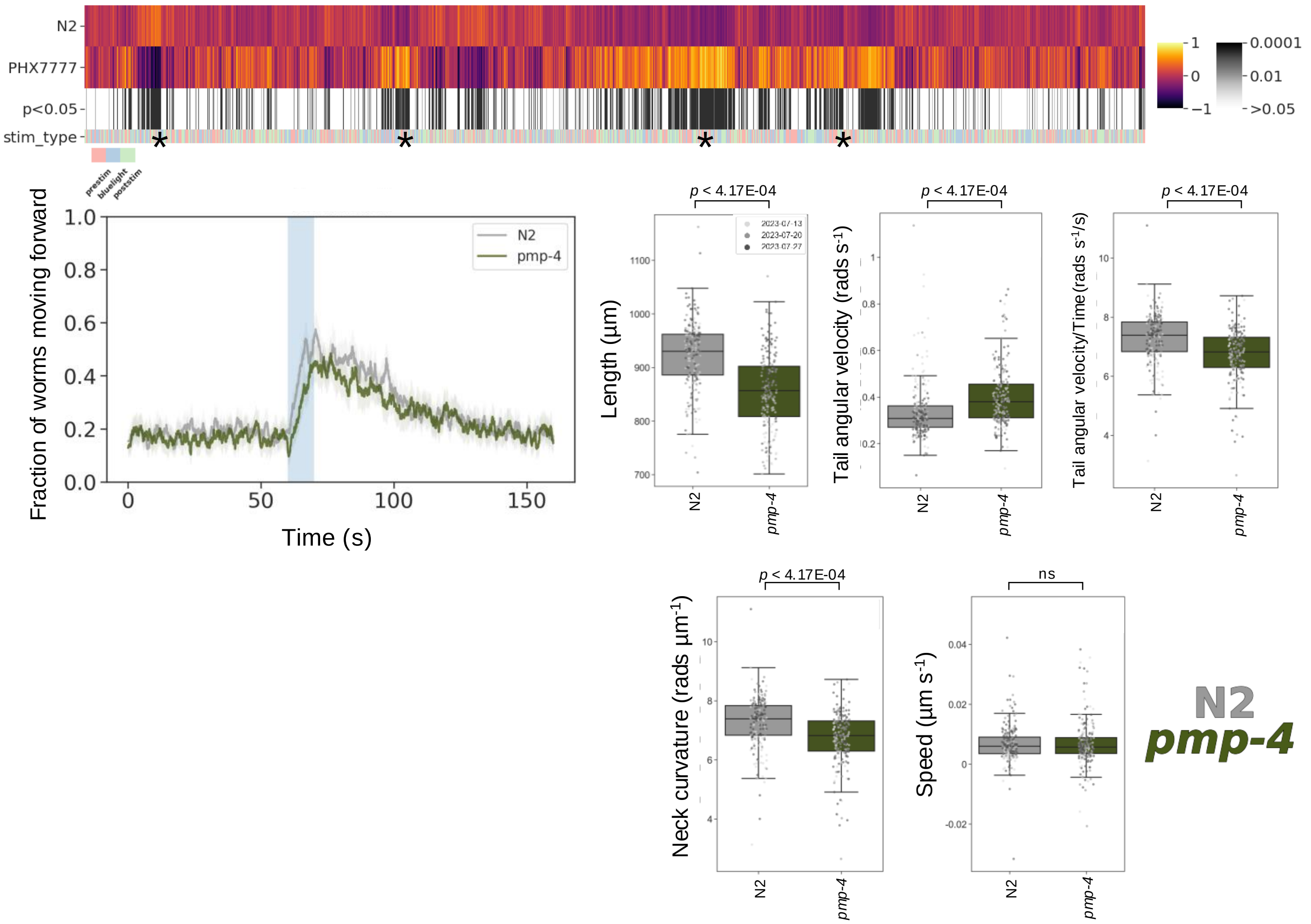

### *R10E11.6(syb7507)*

Strain name: PHX7507
Wild-type gene length: 2858bp
Mutation: 2488bp deletion starting at nt position 26 (deletion of entire gene coding region)

#### *C. elegans* Description

Function: Interacts with several genes, including: *daf-12*, *daf-16* and *gld-1* (function unknown)

Expression: AVK and NSM neurons; anterior hypodermis and germline

Previously reported phenotypes: Neuron degeneration[20]; necrotic cell death variant[21]

#### Human Orthologs:

*SYNRG*

Associated disease(s):
Chromosome 17Q12 Deletion Syndrome; Mixed Phenotype Acute Leukemia

#### Results

5405/8289 significant features vs N2 ($p < 0.05$, block permutation t-test, 10000 permutations)

Key phenotype(s): Shorter; less active, even during stimulation with blue light; decreased angular velocity and curvature (of all body segments); lower frequency of head bends (time derivative of head curvature); and decreased overall speed

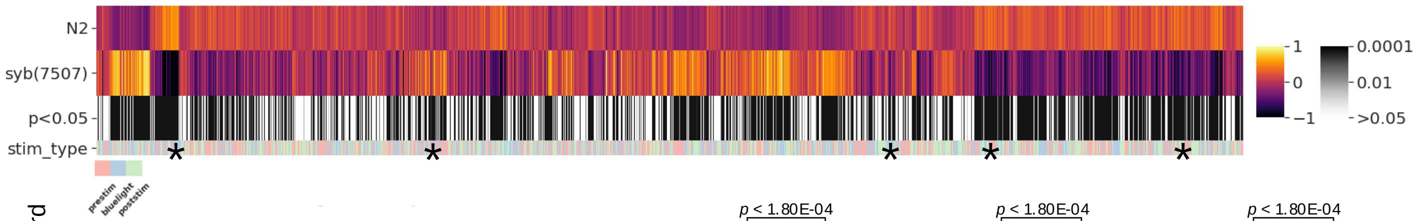


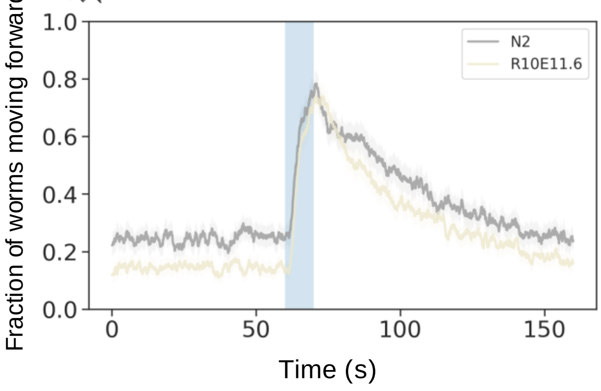


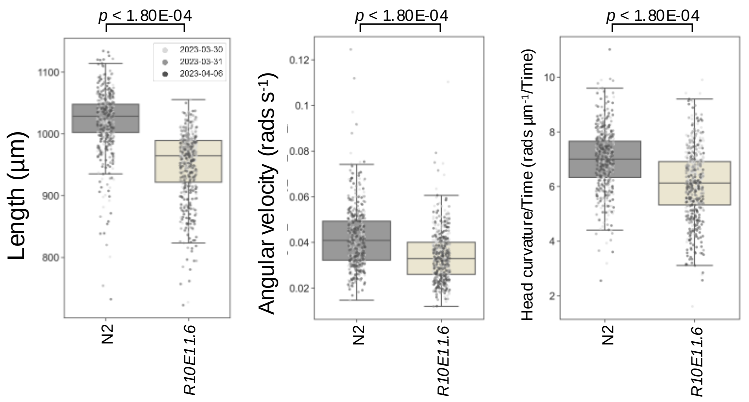


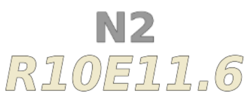


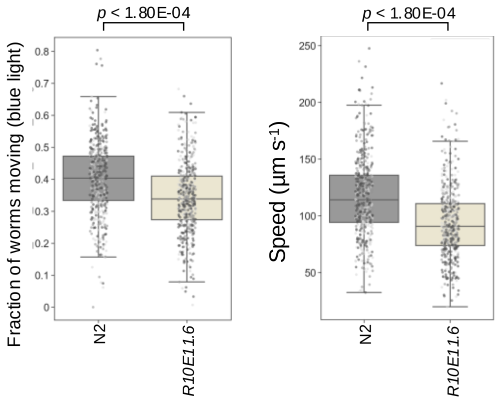

### *rpy-1(syb5027)*

Strain name: PHX5027
Wild-type gene length: 3603bp
Mutation: 3054bp deletion starting at nt position 26 (deletion of entire gene coding region)

#### *C. elegans* Description

Function: Enables acetylcholine receptor binding activity; involved in cholinergic synaptic transmission and positive regulation of neuromuscular synaptic transmission

Expression: Neurons

Previously reported phenotypes: Levamisole resistance[22]; increased fat content[23]

#### Human Orthologs:

*RAPSN*

Associated disease(s):
congenital myasthenic syndrome (11); Fetal akinesia deformation sequence syndrome (2)

#### Results

3661/8289 significant features vs N2 ($p < 0.05$, block permutation t-test, 10000 permutations)

Key phenotype(s): Shorter; decreased angular velocity; increased curvature of midbody and tail; decreased head speed and lower frequency of head bends (time derivative of head curvature); no difference in overall locomotion of the mutant

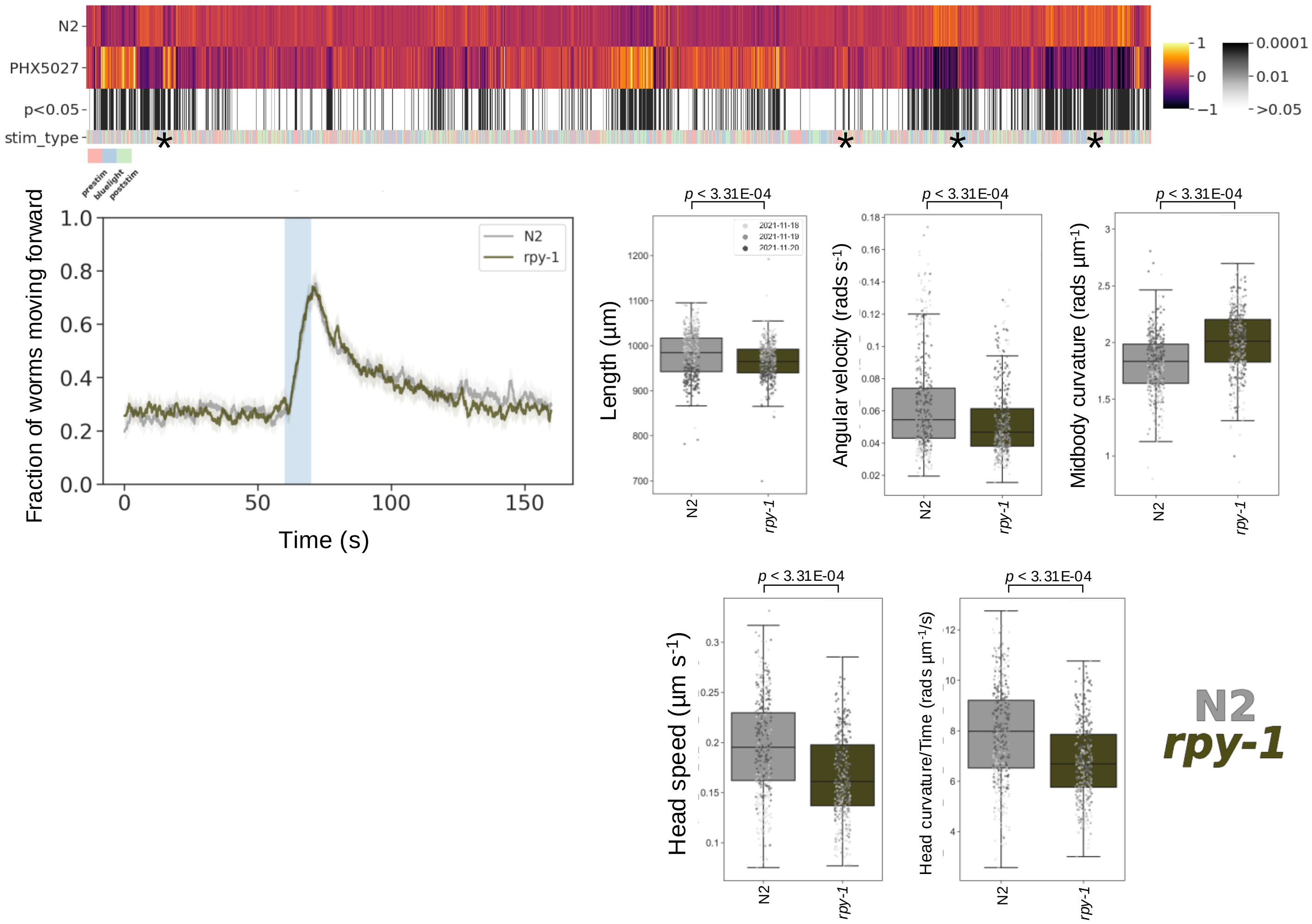

### *sam-4(syb6765)*

Strain name: PHX6765
Wild-type gene length: 1305bp
Mutation: 757bp deletion starting at nt position 234 (half of exon 1 and remaining coding regions)

#### *C. elegans* Description

<u>Function</u>: Enables guanyl-nucleotide exchange factor activity; positive regulator of anterograde synaptic vesical transport (*via* enhancement of UNC-104 recruitment/activation on synaptic vesicles)[26] and pharyngeal pumping[27]

<u>Expression</u>: Nerve ring

<u>Previously reported phenotypes</u>: Dysregulated synaptic vesicle transport[24]; uncoordinated locomotion[25]

#### Human Orthologs:

*BORCS5*

<u>Associated disease(s):</u>
Hermansky-Pudlak syndrome; Colorectal adenocarcinoma; Polymicrogyria; Isolated Corpus Callosum Agenesis

#### Results

3201/8289 significant features vs N2 ($p < 0.05$, block permutation t-test, 10000 permutations)

Key phenotype(s): Shorter; decreased angular velocity, curvature, speed and acceleration of head; no statistically significant difference in other locomotion related features

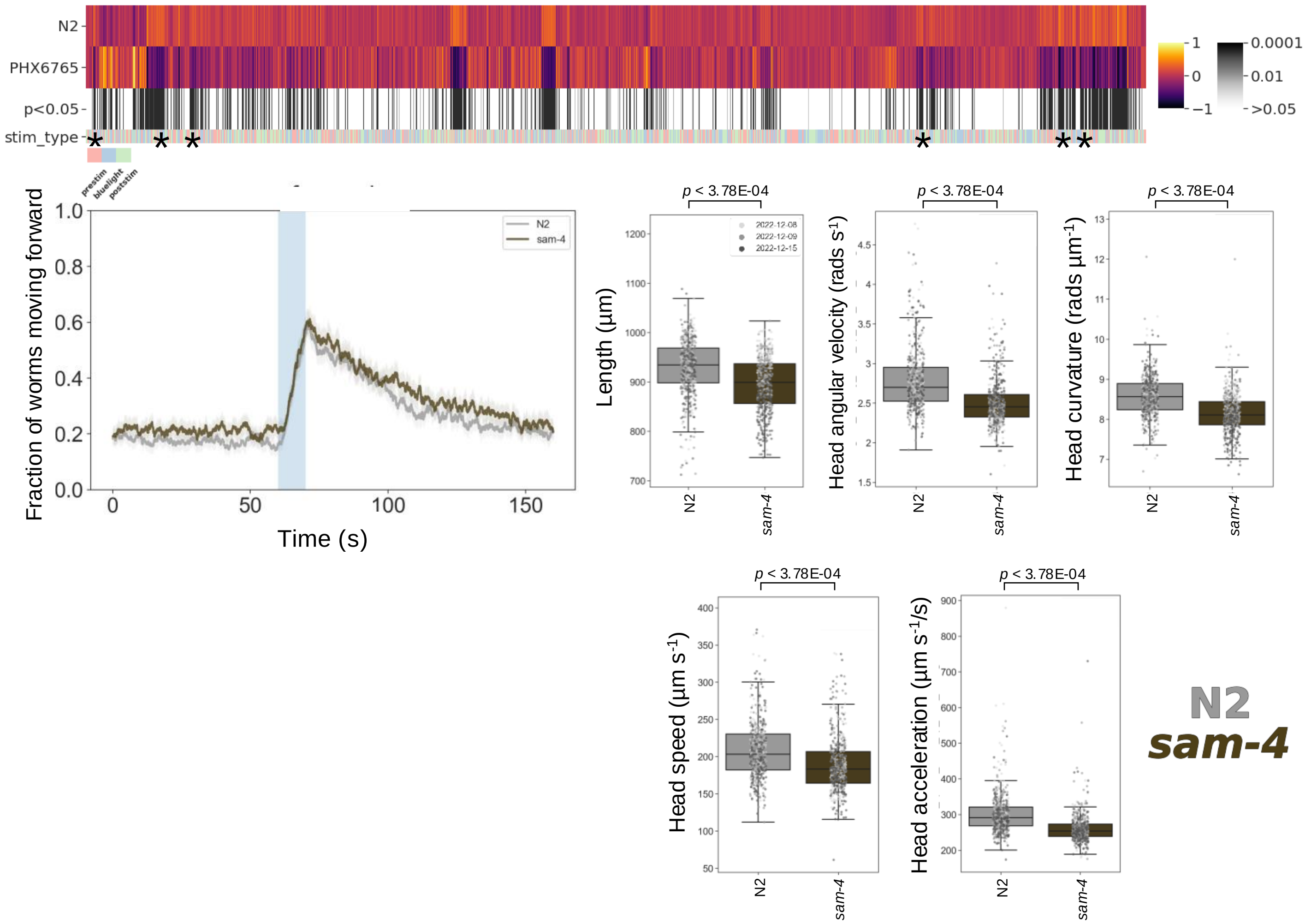

### *sec-31(syb6825)*

Strain name: PHX6825
Wild-type gene length: 5815bp
Mutation: 5815bp deletion starting at nt position 40 (deletion of entire gene coding region)

#### *C. elegans* Description

<u>Function</u>: Predicted to enable structural molecule activity and be involved in COPII-coated vesicle cargo loading, endoplasmic reticulum organisation and intracellular protein transport

<u>Expression</u>: Endoplasmic reticulum exit site

<u>Previously reported phenotypes</u>: Reduced brood size[26]

#### Human Orthologs:

*SEC31A*

<u>Associated disease(s):</u>
Halperin-Birk syndrome; Soft tissue sarcoma

#### Results

4236/8289 significant features vs N2 ($p < 0.05$, block permutation t-test, 10000 permutations)

Key phenotype(s): Longer; decreased angular velocity of head and tail; decreased curvature of head and neck; lower frequency of midbody bends (time derivative of midbody curvature); no difference in locomotion features

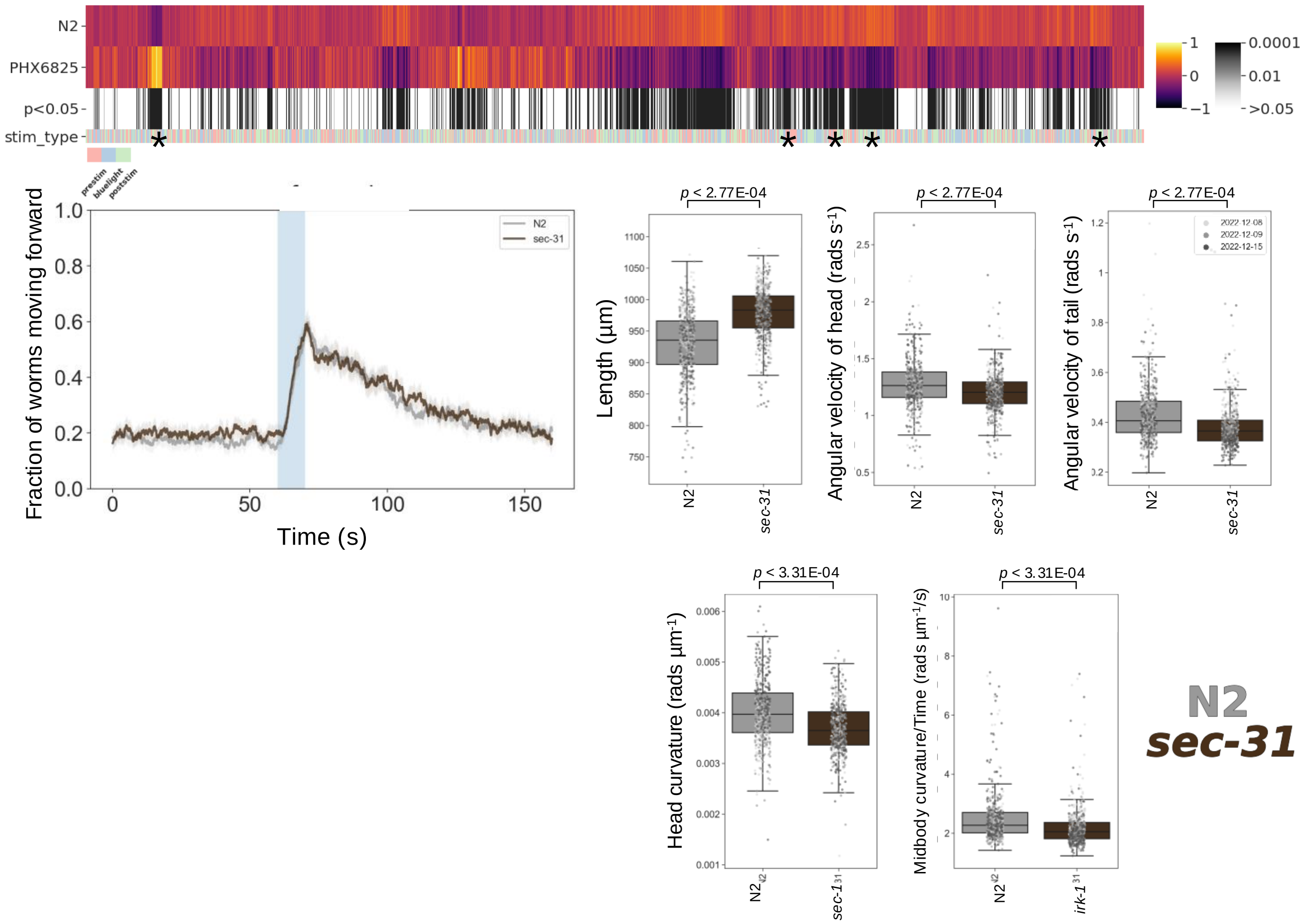

### *shl-1(syb5907)*

Strain name: PHX5907
Wild-type gene length: 5589bp
Mutation: 5166bp deletion starting at nt position 91 (start of exon 1 to end of exon 5)

#### *C. elegans* Description

Function: Predicted to enable transient outward potassium channel activity; involved in potassium ion transmembrane transport

Expression: Body wall musculature; muscle cell neurons; pharynx; tail

Previously reported phenotypes: Reduced male mating efficiency[27]; reduction of peak transient (electrical) current of myocytes[28]

#### Human Orthologs:

*KCND2* and *KCND3*

Associated disease(s):
Brugada syndrome (9); Spinocerebellar ataxia (types 19 and 22)

#### Results

289/8289 significant features vs N2 ($p < 0.05$, block permutation t-test, 10000 permutations)

Key phenotype(s): Fatter; decreased angular velocity of head while moving; decreased head acceleration (time derivative of head speed); attenuated backing, but sustained forward response, upon stimulation with blue light

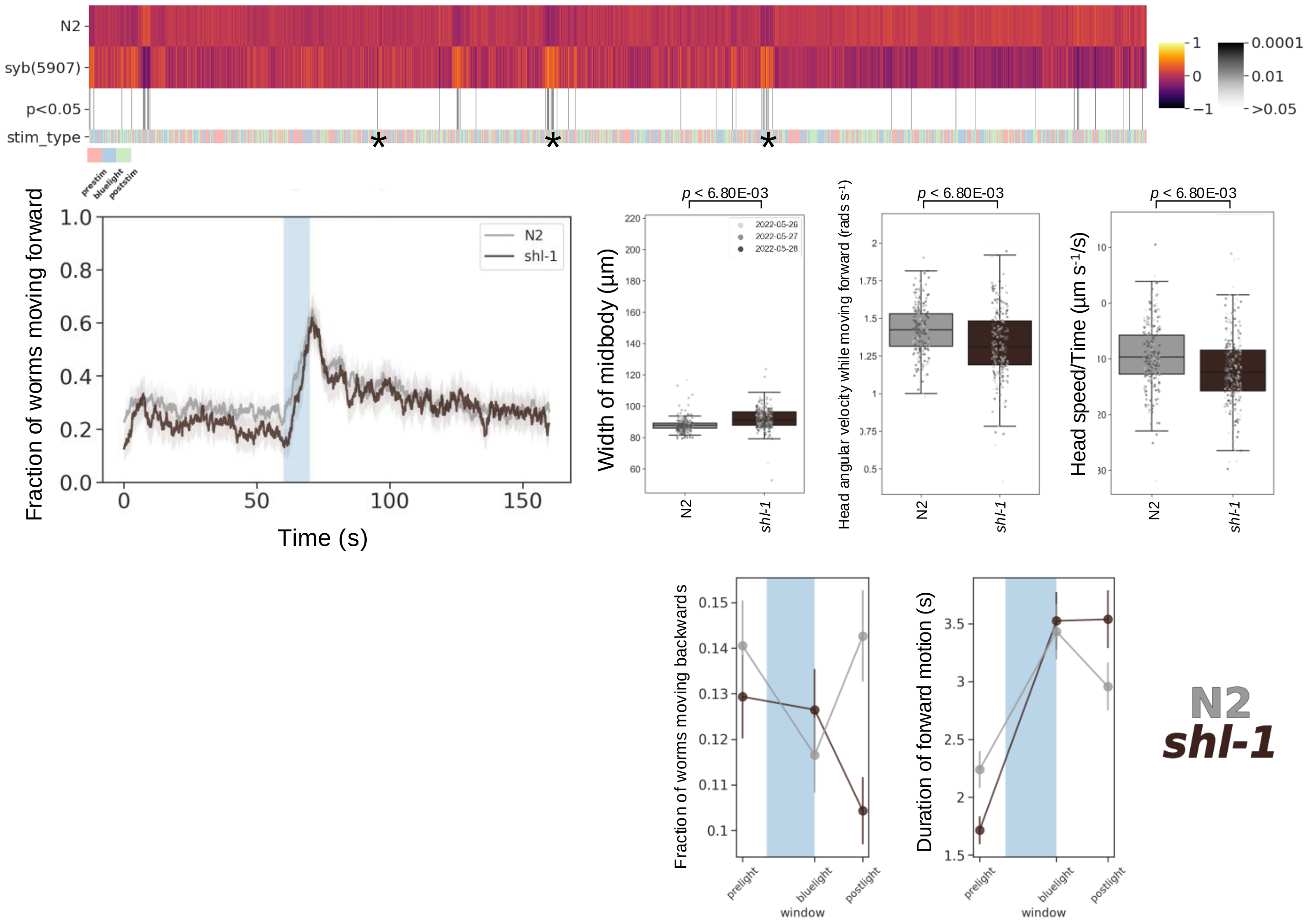

### ***smc-3(fq180)***

Strain name: *smc-3*((*fq180*)[K115E] III)
Wild-type gene length: 4003bp
Mutation: K to E amino acid substitution position 115. Synonymous to K114E mutation in human *SMC3*

#### *C. elegans* Description

Function: Enables cohesion loader activity and double-stranded DNA binding activity; involved in mitotic sister chromatid cohesion

Expression: Chromatin and cohesion complex

Previously reported phenotypes: Embryonic lethality (for genetic knockdown)[29]; reduced brood size[30]; meiosis variant[30]

#### Human Orthologs:

*SMC3*

Associated disease(s):
Cornelia de Lange syndrome; Hepatocellular carcinoma; Intellectual disability

#### Results

5685/8289 significant features vs N2 ($p < 0.05$, block permutation t-test, 10000 permutations)

Key phenotype(s): Longer; increased midbody curvature; decreased curvature and angular velocity of the head; less active during baseline tracking, with no difference in photophobic escape response; increased duration of pausing following stimulation with blue light

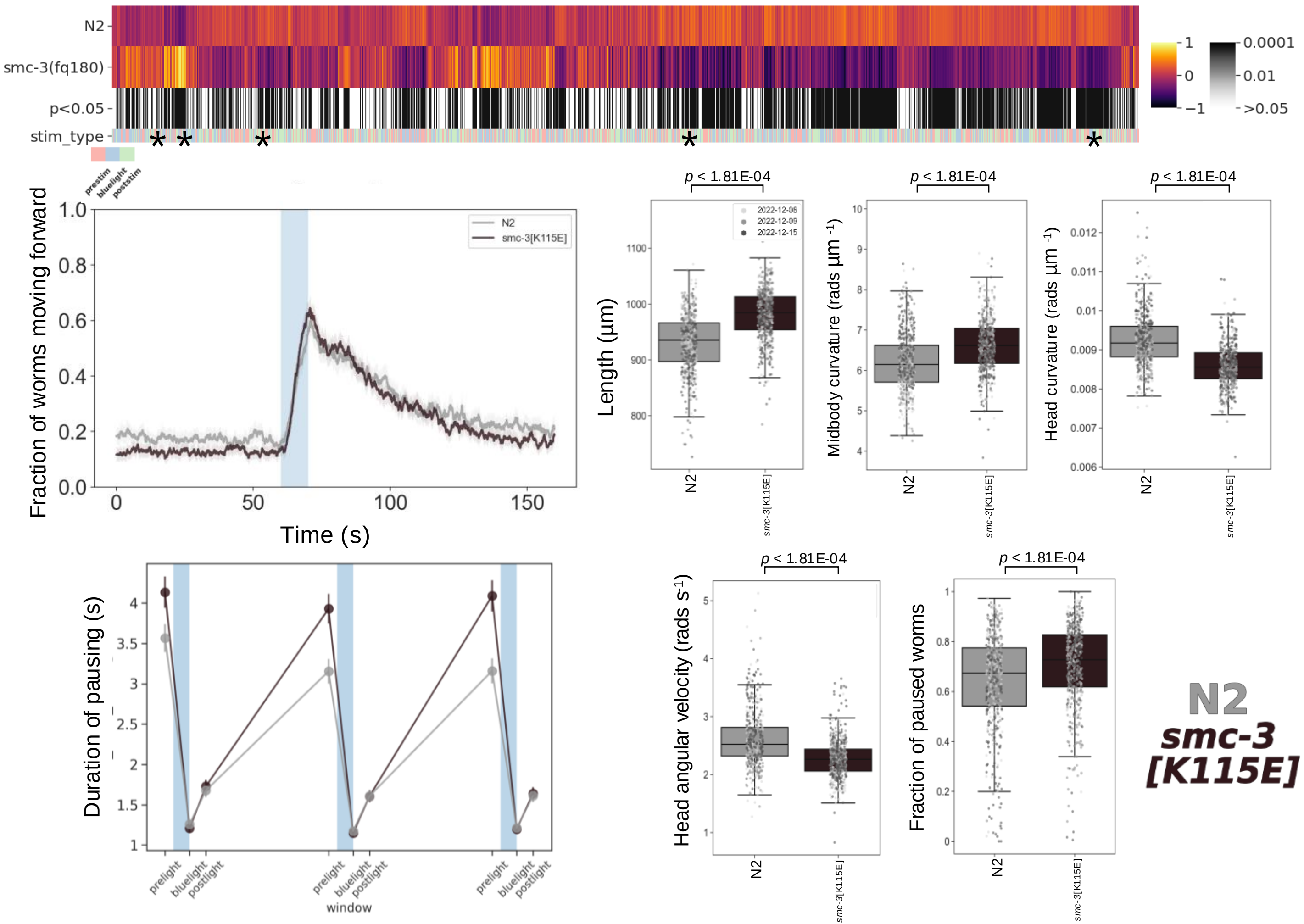

### ***tmem-222(syb4882)***

Strain name: PHX4882
Wild-type gene length: 657bp
Mutation: 544bp deletion starting at nt position 16 (deletion of entire gene coding region)

#### *C. elegans* Description

Function: Predicted to be an integral membrane component

Expression: Uncharacterised

Previously reported phenotypes: None

#### Human Orthologs:

*TMEM222*

Associated disease(s):
Neurodevelopmental disorder with motor and speech delay; Non-specific syndromic intellectual disability

#### Results

68/8289 significant features vs N2 ($p < 0.05$, block permutation t-test, 10000 permutations)

Key phenotype(s): Increased midbody curvature

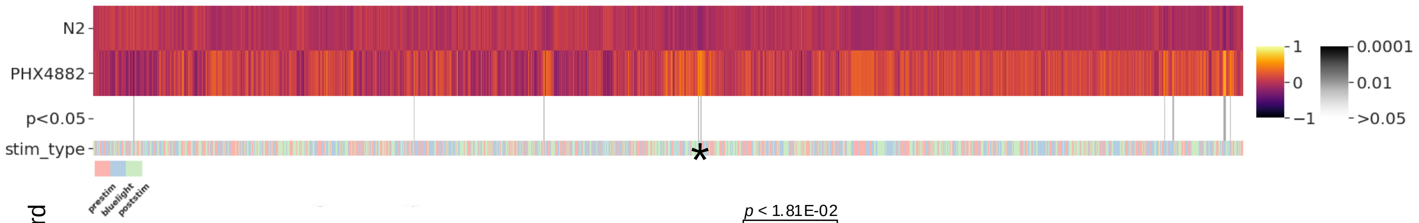


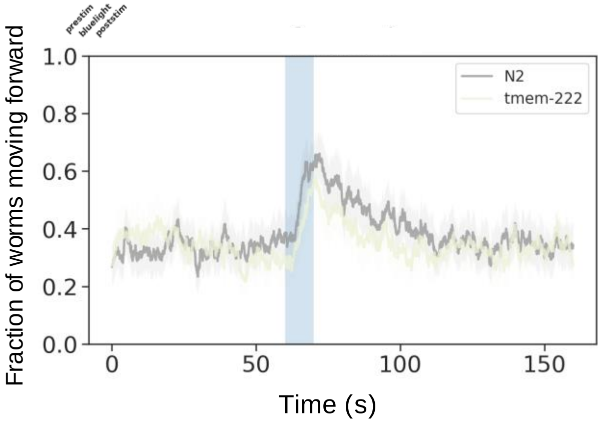


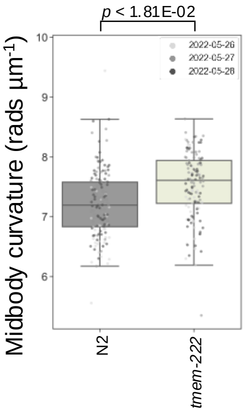


**N2**
***tmem-222***

### *vps-50(syb6653)*

Strain name: PHX6653
Wild-type gene length: 5137bp
Mutation: 5111bp deletion starting at nt position 201 (deletion of entire gene coding region)

#### *C. elegans* Description

Function: Predicted to enable SNARE binding activity; positive regulation of dense core granule transport; regulation of locomotory behaviour

Expression: Cytosol and EARP complex

Previously reported phenotypes: Increased duration of egg retention[31]; lack of slowing in response to food[31]

#### Human Orthologs:

*VPS50*

Associated disease(s):
Neurodevelopmental disorder with microcephaly; Seizures; Neonatal cholestasis; Pontocerebellar hypoplasia, Type 2E

#### Results

6952/8289 significant features vs N2 (*p* < 0.05, block permutation t-test, 10000 permutations)

Key phenotype(s): Less active during baseline recordings; strong, but short-lived (increased paused duration), forward photophobic escape response; sustained backward escape response; shorter; decreased angular velocity and curvature (for all body segments); slower

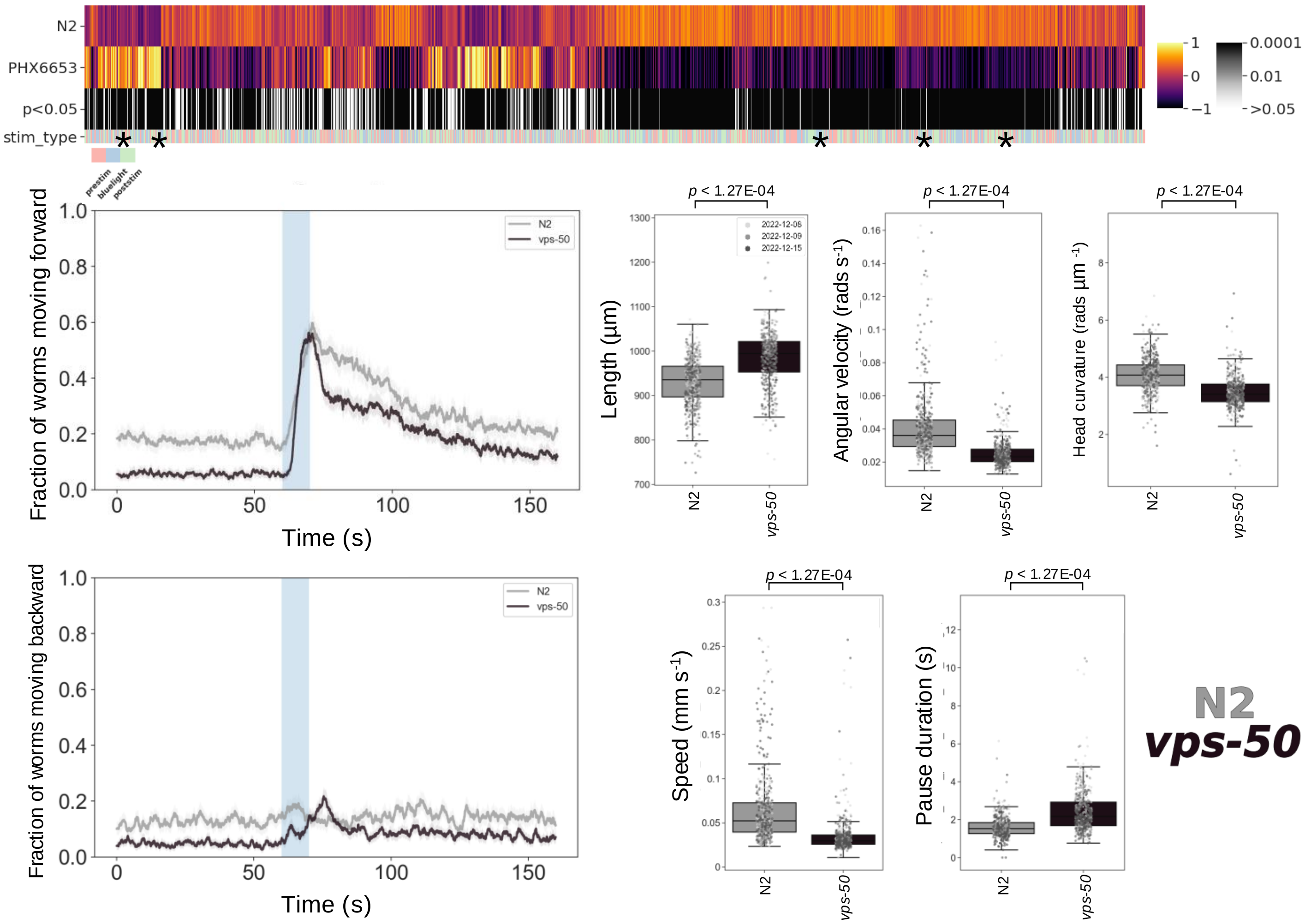

### *Y47D9A.1(syb7612)*

Strain name: PHX7612 *Y47D9A.*1(*syb7612*)[R298W]
Wild-type gene length: 983bp
Mutation: R to W amino acid substitution position 298- Synonymous to R318W mutation inhuman GMPPA

#### *C. elegans* Description

Function: Predicted to enable transferase activity

Expression: Cytoplasm (predicated)

Previously reported phenotypes: None

#### Human Orthologs:

*GMPPA*

Associated disease(s):
Alacrimia; Achalasia; Impaired intellectual development syndrome

#### Results

3704/8289 significant features vs N2 ($p < 0.05$, block permutation t-test, 10000 permutations)

Key phenotype(s): Attenuated photophobic escape response; slow moving; shorter; decreased angular velocity; increased frequency of body bends (time derivative of midbody curvature)

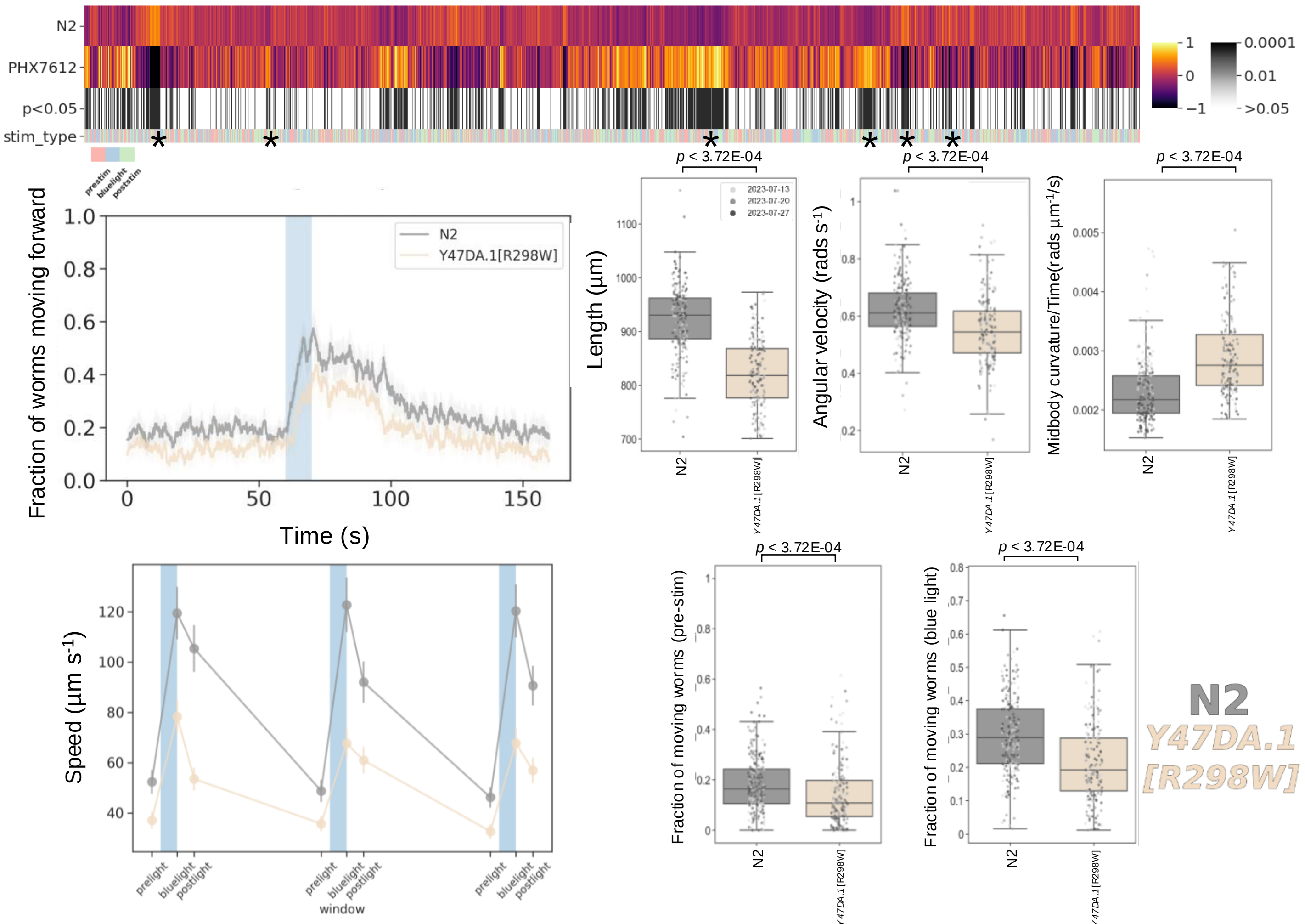
